# Resequencing 250 soybean accessions: new insights into genes associated with agronomic traits and genetic networks

**DOI:** 10.1101/2021.01.28.428693

**Authors:** Chunming Yang, Jun Yan, Shuqin Jiang, Xia Li, Haowei Min, Xiangfeng Wang, Dongyun Hao

## Abstract

Limited knowledge on genomic diversity and the functional genes associated with soybean variety traits has resulted in slow breeding progress. We sequenced the genome of 250 soybean landraces and cultivars from China, America and Europe, and investigated their population structure, genetic diversity and architecture and selective sweep regions of accessions. We identified five novel agronomically important genes and studied the effects of functional mutations in respective genes. We found candidate genes *GSTT1*, *GL3* and *GSTL3* associated with isoflavone content, *CKX3* associated with yield traits, and *CYP85A2* associated with both architecture and yield traits. Our phenotype-gene network analysis revealed that hub nodes play a role in complex phenotypic associations. In this work, we describe novel agronomic trait associated genes and a complex genetic network, providing a valuable resource for future soybean molecular breeding.

## Introduction

Soybean *Glycine max* [L.] Merr. is one of the most important crops worldwide of vegetable oil and proteins source for human and livestock feed etc. Soybean originated in China and its wild species (*G. soja* Sieb. & Zucc.) were domesticated in approximately 3,000 B.C. before introduced to Korea and Japan about 3,000 years later. It was brought to Europe and North America in the 18th century, and extensively cultivated on a global scale since the 19th century[1].

With the rapid development of modern molecular biology and the high-throughput sequencing technologies, whole-genome resequencing and genome wide association studies (GWAS) have become common methods used to study population genetic diversity and locating phenotypic related quantitative trait loci (QTL) or genes. This has improved our knowledge extensively in crop genomes and selective breeding. In recent years, for example, there are increasing number of reports on the domestication and improvement of soybean at the genome-wide level. This includes genes and genetic networks related to soybean agronomic traits and functions[2–4]. However, due to the diversity of soybean varieties and their complex genetic background, our knowledge of the soybean genome and functional genes is still limited in comparison to rice and maize. A greater number of soybean varieties need further exploration at the genomic level, particularly in relation to molecular traits associated with edible quality, soybean “ideotype” and the underlying genetic network of high-yielding varieties.

In this study, we collected 250 soybean varieties from the core Northeast China soybean germplasm pool, which consisted of 134 accessions of landrace and cultivar from Northeast and Northwest China, as well as 116 accessions from European and North American cultivars. The genomes of the most accessions are not sequenced previously. We performed the high-depth whole-genome resequencing and comprehensive analyses of this 250 soybean population. The generated dataset revealed valuable new information on soybean genome structure, novel genes associated with important agronomic traits and the genetic networks. These genetic resources provide unique references into molecular breeding and evolution study in soybean.

## Results

### Genome resequencing and variation calling

High-depth whole genome resequencing was performed on 250 soybean accessions, including 51 landraces and 83 cultivars originating from provinces in Northeast China (i.e. Heilongjiang, Jilin, Liaoning and Northeast Inner Mongolia) and Northwest China (i.e. Xinjiang, Ningxia and Gansu), as well as 116 cultivars originating from Europe and North America **(Figure 1a, Supplementary Table 3).** In total, we obtained approximately 10G of pair-end reads and 3T bases. The maximum sequencing depth of a single accession was 22.5x, with the average depth at 11x. After filtering out the raw sequencing data (see methods), the remaining high-quality cleaned data were compared with the soybean reference genome *G. max v2.0[5].* The effective mapping rates ranged from 74.8% to 87.6%, while the genome coverage ranged from 94.8% to 97.0% **(Supplementary Table 1)**. The high mapping rates and coverage guarantee that the sequenced data is reliable and of high quality.

**Figure 1.**
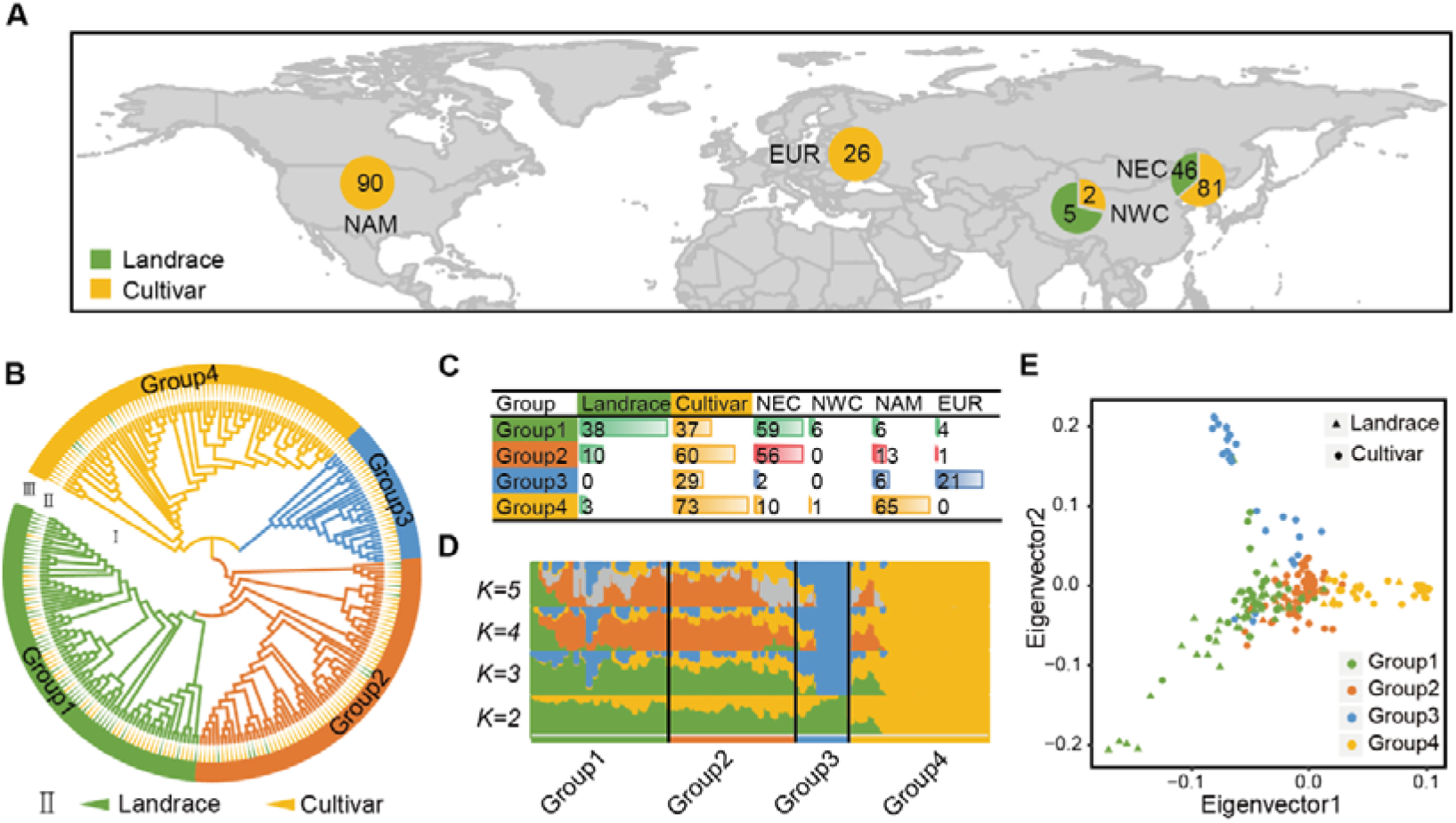
Population structure of 250 soybean accessions. **A.** Geographic distribution of 250 soybean landraces and cultivars. Landraces are shown with green color and cultivars are shown with yellow color. **B.** Phylogenetic tree constructed for all soybean accessions. Group1-4 are shown with different colors, Landraces are labeled with green triangles and cultivars are labeled with yellow triangles. **C.** Statistics of the geographic origin for each subpopulation. **D.** Mixed ancestors analysis for soybean subpopulations. Each color represents an ancestral component. K from 2 to 5 are set to trace different ancestral components. **E.** PCA plot of the first two eigenvectors for all soybean accessions. Landraces and cultivars are shown will different shape, while groups are shown with different colors.

Through standard variation detection, genotype filtering and imputation steps (see methods), we detected in total 6,333,721 single nucleotide polymorphisms (SNPs) and 2,565,797 insertion & deletions (Indels). This includes 244,360 SNPs and 62,714 Indels located in the exon regions. The ratio of non-synonymous SNP to synonymous SNP substitution was 1.37. There are 4,311,814 SNPs with a minor allele frequency (MAF) larger than 0.05 **(Supplementary Table 2 & Supplementary Figure 1**). In summary, we achieved over 6M high-density and high-quality genotype data from 250 soybean accessions with a density of one SNP per 15 bases.

### Population structure of soybean landraces and cultivars

Using the 6M SNP genotype dataset, we constructed a phylogenetic tree using the neighbour joining (NJ) method. This resulted in the classification of the 250 soybean accessions into four groups (**Figure 1b**). Among them, *Group 1* included 65 Chinese varieties, four European and six American cultivars, whereas *Group 2* contained 56 Chinese varieties, one European and 13 American varieties. In *Group 3*, there were 21 European, two Chinese and six American cultivars, while 65 North American cultivars and 11 Chinese varieties were clustered within *Group 4* (**Figure 1c**). Principal Component Analysis (PCA) results were consistent with the phylogenetic tree results. Three groups, *Group 1*, *3*, and *4* radiated away from *Group 2* within the rectangular coordinate system projected using eigenvector 1 and eigenvector 2 data on X and Y axes, respectively. Concurrently, the distribution of varieties in the four groups had continuity, indicating varieties located in different groups also have genetic similarities **(Figure 1e**). A Bayesian clustering algorithm based on a mixed model was used to estimate the proportion of ancestors in each accession. That is, when *K = 2*, the main ancestor component (yellow) of *Group 4* was split, indicating that *Group 4* has the highest level of selection. When *K = 3*, the main ancestor component (blue) of *Group 3* was split, indicating that *Group 3* has the second level of selection. However, when *K = 4* and *K = 5*, *Group 1 and Group 2* exhibit complex differentiated mixed ancestor components, indicating a higher genetic diversity and lower selection level in *Groups 1* and *2* (**Supplementary Table 3, Figure 1d**).

These results indicate that group classification of the 250 soybean accessions is closely related to their geographical distribution. That is, varieties with similar geographical distribution have similar genetic backgrounds. Generally speaking, the group classification was also related to the level of domestication. Landraces have a lower level of domestication, while cultivars have higher levels of domestication. Varieties with similar domestication levels tend to have a higher similarity in genetic backgrounds. However, there are still differences in geographical distribution and domestication level among breeds with similar genetic backgrounds, indicating that gene exchange may have occurred between accessions of different groups. This observation reflected the complexity of soybean domestication history.

### Genetic diversity and selective sweep analysis

Linkage disequilibrium (LD) analysis showed that the overall LD decay distance was more than 100 kb, and the LD decay distance of the landraces was smaller than that of the cultivars (**Figure 2A**). Further LD decay analysis of the four groups showed that the LD decay distance of *Group 1* was the smallest, followed by *Groups 2* and *3*, while *Group 4* had the largest LD decay distance (**Figure 2B**). In addition, the LD levels varied for different chromosomes or different regions across one chromosome. Identical by state (IBS) analysis can reflect the degree of relatedness among individuals by calculating the consistency of all genetic markers. The IBS values of all comparisons in each group were calculated, and it was found that the average IBS values of landraces were less than that of cultivars (**Figure 2C**). The IBS values of *Groups 1*-*4* followed the same trends as that of LD decay distance. In particular, the IBS values of *Group 1* were the lowest and the IBS values of *Group 4* were the highest of all groups (**Figure 2D**). *θπ* values can reflect the genetic diversity within a population by calculating the number of different sites between any two sequences or individuals within a population. *Fst* is a calculation used to measure the differentiation and genetic distance between two populations. *θπ* values were calculated for landraces, cultivars, all accessions, and *Groups 1*-*4*. *Fst* values were calculated between landraces and cultivars, and between each comparison of the four Groups. Results show that a population with a higher level of LD decay distance or higher IBS values correlate with a smaller *θπ* (**Figure 2E, F**). This pattern is opposite to that of the LD decay and IBS values. The lowest *Fst* value was for *Group 1* versus *Group 2*, while the highest value was for *Group 3* versus *Group 4*. We also observed that the *Fst* value of *Group 2* versus *Group 3* was higher than that of *Group 1* versus *Group 3* (**Figure 2G**). The *Fst* value of *Group 2* versus *Group 4* was smaller than that of *Group 1* versus *Group 4*. In addition, the results of allele frequency distribution (AFD) analysis, as an alternative population similarity measurement, were consistent with the *Fst* results (**Supplementary Figure 2**). The population diversity analysis results, when combined with population structure and geographical distribution information, infer that the European and American soybean varieties may have originated from different Chinese ancestors before undergoing independent selection. Results indicate that European cultivars and the Chinese landrace group (*Group 1*) in our study have a more recent common ancestor, while North American cultivars and the Chinese cultivar group (*Group 2*) have a more recent common ancestor.

**Figure 2.**
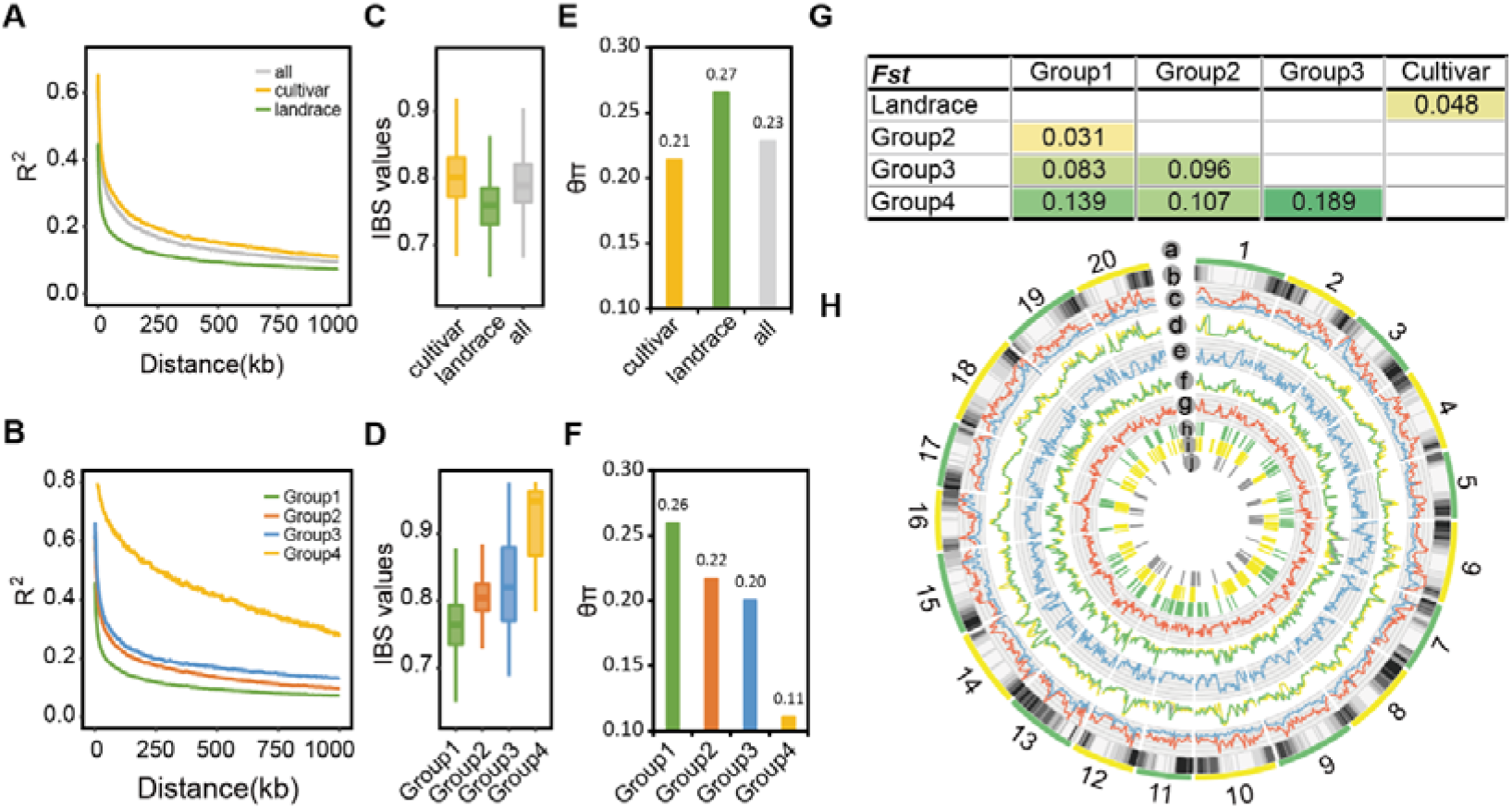
Genetic diversity of soybean subpopulations. **A.** LD decay plots for landrace (green), cultivars (yellow) and all soybean accessions (grey). **B.** LD decay plots for soybean subpopulations. **C.** IBS values distribution for landrace (green), cultivars (yellow) and all soybean accessions (grey). **D.** IBS values distribution for soybean subpopulations.**E.** Comparison of *θπ* values for landrace (green), cultivars (yellow) and all soybean accessions (grey). **F.** Comparison of *θπ* values for soybean subpopulations. **G.** Comparison of *Fst* values between subpopulations. **H.** Landscape of soybean genetic diversity across the whole genome. (a) Chromosomes. (b) Density of genes (c) Density of SNPs (red) and Indels (blue). (d) LD values distribution for landraces(green), cultivars(yellow) and all accessions(grey). (e) *Fst* values distribution of landraces versus cultivars (f) *θπ* values distribution for landraces(green), cultivars(yellow) and all accessions(grey). (g) *Tajima’D* values distribution of all accessions. (h) Putative selective sweep regions detected by *Tajima’D* combine *θπ*. (i) Putative selective sweep regions detected by *Fst* combine *θπ* ratios. (j) ROH region larger than 300 Kb.

*Tajima’ D* (based on a neutral test), *θπ* (based on genetic diversity within a population), and *Fst* (based on genetic diversity between two populations), have provided us with highly effective tools that screen selective sweep signals across a genome[6]. We combined methods in pairs for mining potential selective sweep regions in the soybean genome that may have underwent artificial selection. One pair was *Tajima’ D* combined with *θπ* for the whole population. Another pair was *Fst* combined with *θπ* ratios between two subpopulations, landrace and cultivar. We used a sliding window method to calculate the values of *Tajima’ D, θπ, and Fst* in each window across the whole genome, and selected the top 5% most significant windows as potential selective sweep regions (**Supplementary Figure 3A, B**). A total of 148 and 222 potential selective sweep regions were screened by the two methods, and they covered 36.09 Mb and 88.15 Mb genome regions, respectively (**Supplementary Table 4**). These potential selective sweep regions covered 9,128 genes, accounting for approximately one sixth of all soybean genes. A total of 1,876 genes were screened by both methods (**Supplementary Figure 3C**). A runs of homozygosity (ROH) region is a continuous homozygous chromosome region in a genome, which may relate to domestication or artificial selection[7]. Through ROH analysis, 71 ROH regions larger than 300 kb were obtained from all 250 accessions, with a total length of 27.84 Mb. The longest ROH regions up to 911 kb were located at the beginning of chromosome 10 (**Supplementary Table 5**). There were 3,397 genes located in these ROH regions, 924 of which were also located in the potential selected sweep region (**Supplementary Figure 3D**).

### Phenotype-related loci and genes identified using GWAS analysis

We measured 50 agronomic traits in 250 soybean accessions from three geographic locations for three years, and then integrated them using best linear unbiased prediction (BLUP). The 50 traits included traits related to architecture (15), colour (5), isoflavone (1), oil (4), protein (18) and yield (7) and were classified into six categories (**Supplementary Table 6**). We calculated Pearson correlation coefficients for traits so as to compare within and between categories, and found that traits in the same categories were more strongly correlated than traits in different categories. For example, there were strong positive or negative correlations between almost all protein-related traits, oil-related traits and yield-related traits. Linoleic acid content was positively correlated with linolenic acid content, but negatively correlated with oleic acid content. Stem intension was negatively correlated with lodging (**Supplementary Figure 4**). Some traits were evenly distributed, while others were ranked (**Supplementary Figure 5-54**).

Using 4,311,814 SNPs with a MAF > 0.05 as an input, we performed GWAS analysis using the mixed linear model (MLM) method for 50 agronomic traits. For each trait, we used a clump based method[8] and defined a significant associated loci (SAL) at a chromosome region with a substantial amount of SNPs associated with the trait. A total of 203 SALs were detected in 43 traits (**Supplementary Figure 5-54, Supplementary Table 7**). Since each SAL may contain dozens of genes, we used a functional mutation based haplotype test method for further mining of the most reliable candidate trait associated genes[9]. In particular, we considered only the non-synonymous SNPs, frameshift Indels, mutations within a gene that happened on a start or stop codon, splice site or transcription start sites as effective functional mutations. We used these mutations to classify a gene into different haplotypes, and subsequently tested the phenotypic differences of the accessions belonging to each haplotype. A gene with significant phenotypic differences was defined as significant associated gene (SAG), and 3,165 SAGs were screened in 43 traits. These SAGs include some QTL or genes that have been previously identified, such as: the flower colour related *chr13:16551728-19506795*; pubescence colour related *chr6:16930159-19168772*; seed coat lustre related *chr15:8910798-10281804*; palmitic acid content related *chr5:879095-1682551[4]*; isoflavone content related *chr5:38880530-39142565[10]*; plant height related *Dt1[4]*; and oil content related *FAD2* and *SAT1[11]*, among others. They also contain genes that we have identified for the first time in soybean, such as: the isoflavone content related *GL3* and *GSTL3*; the yield traits related *CKX3*; and the architecture and yield traits related *CYP85A2*.

### Novel genes related to isoflavone content

Isoflavone content is an important quality-related trait in soybean, but its molecular mechanism is still unclear. Here we identified four SALs related to isoflavone content, namely *chr3:38590023-38728718*, *chr5:3888053-39142565*, *chr13:18342836-18541809*, and *chr5:24726091-24852447*. Only one SAL *chr5:24726091-24852447* overlaps with a previously reported QTL that contains a *GST* (Glutathione S-transferase) gene *GSTT1*[10]. All other SALs are newly identified. There are 48 genes located within these SALs (**Supplementary Table 8**), and three genes (*GSTT1, GL3* and *GSTL3)* may be related to isoflavone content (**Figure 3A, B**). There are two functional mutation sites at *c5s38936266* and *c5s38940717*, forming two haplotypes for *GSTT1a* and *GSTT1b*, respectively. For each *GSTT1* gene, soybean accessions with a different haplotype have significantly different isoflavone contents. Since *GSTT1a* and *GSTT1b* are approximately only 1 kb apart from each other in the same genome region, we considered the two genes to be one in further analysis. Three haplotypes were formed by two functional mutation sites when the two *GSTT1* genes were analysed as one. *Haplotype 1* versus *Haplotype 2*, as well as *Haplotype 2* versus *Haplotype 3* showed significant differences in isoflavone content, while *Haplotype 1* versus *Haplotype 3* showed no significant differences (analysed using Tukey’s test). This suggests that *GSTT1a* is associated with isoflavone content due to its linkage with *GSTT1b*. However, *c5s38936266* did not contribute to the isoflavone content difference. Thus, only *c5s38940717* on *GSTT1b* was associated with isoflavone content (**Figure 3C**).

**Figure 3.**
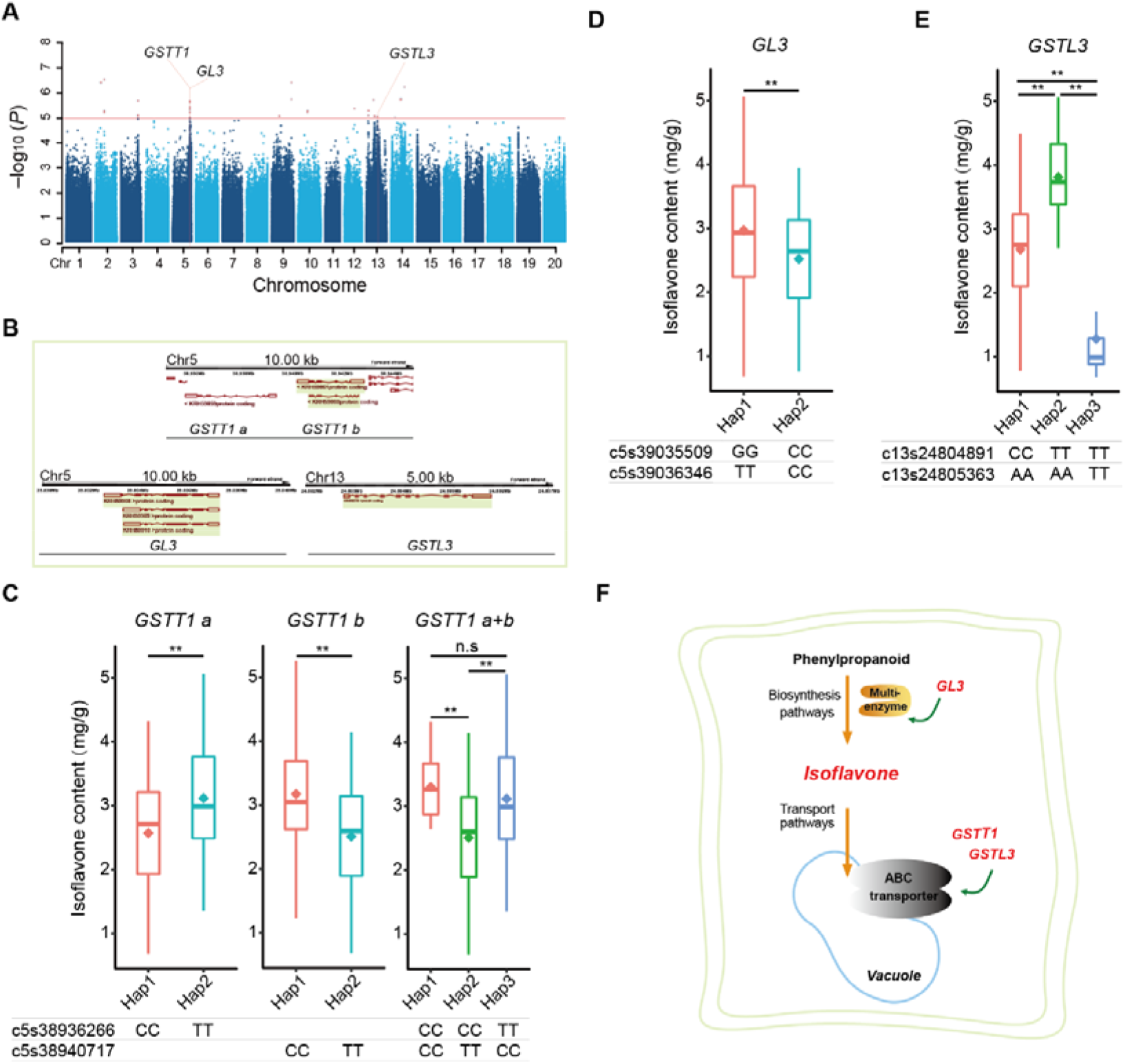
GWAS of soybean isoflavone content. **A.** Manhantan plot and four candidate genes for soybean isoflavone content. **B.**Chromosome location and transcripts structure of the candidate genes. **C.** Soybean isoflavone content distribution for the haplotypes of gene *GSTT1*. **D.** Soybean isoflavone content distribution for the haplotypes of gene *GL3*. **E.** Soybean isoflavone content distribution for the haplotypes of gene *GSTL3*. **F.** Diagram of soybean isoflavone synthesis and transport, and the roles of candidate genes detected by GWAS. (\**P* < 0.05; \*\**P* < 0.01; n.s., not significant)

We found that two functional mutations, *c5s39035509* and *c5s39036346* producing two haplotypes in *GL3*, were associated with different isoflavone contents in soybean accessions (**Figure 3D**). We also identified another *GST* gene, *GSTL3,* which was located in chromosome 13. Two functional mutations produced three haplotypes, and significant associations between the different haplotypes and isoflavone content was detected for each comparison (**Figure 3E**). Based on the above results, we drew a schematic diagram of the roles of candidate genes according to their biological functions where we indicate that *GL3* regulates isoflavone synthesis, whilst *GSTT1* and *GSTL3* participate in isoflavone transport (**Figure 3F**).

### Yield related traits and the artificial selection of CKX3

Four yield related traits (pod number per plant, seed number per plant, one hundred seed weight and seed size) have a common SAL located at the ~4.0 Mb to ~4.2 Mb region of chromosome 17 (**Figure 4A**). Further analysis revealed that this SAL contains two tandem repeat *CKX* (cytokinin oxidase/dehydrogenase) genes named *CKX3* and *CKX4*, approximately 15 kb apart from each other (**Figure 4F**).

**Figure 4.**
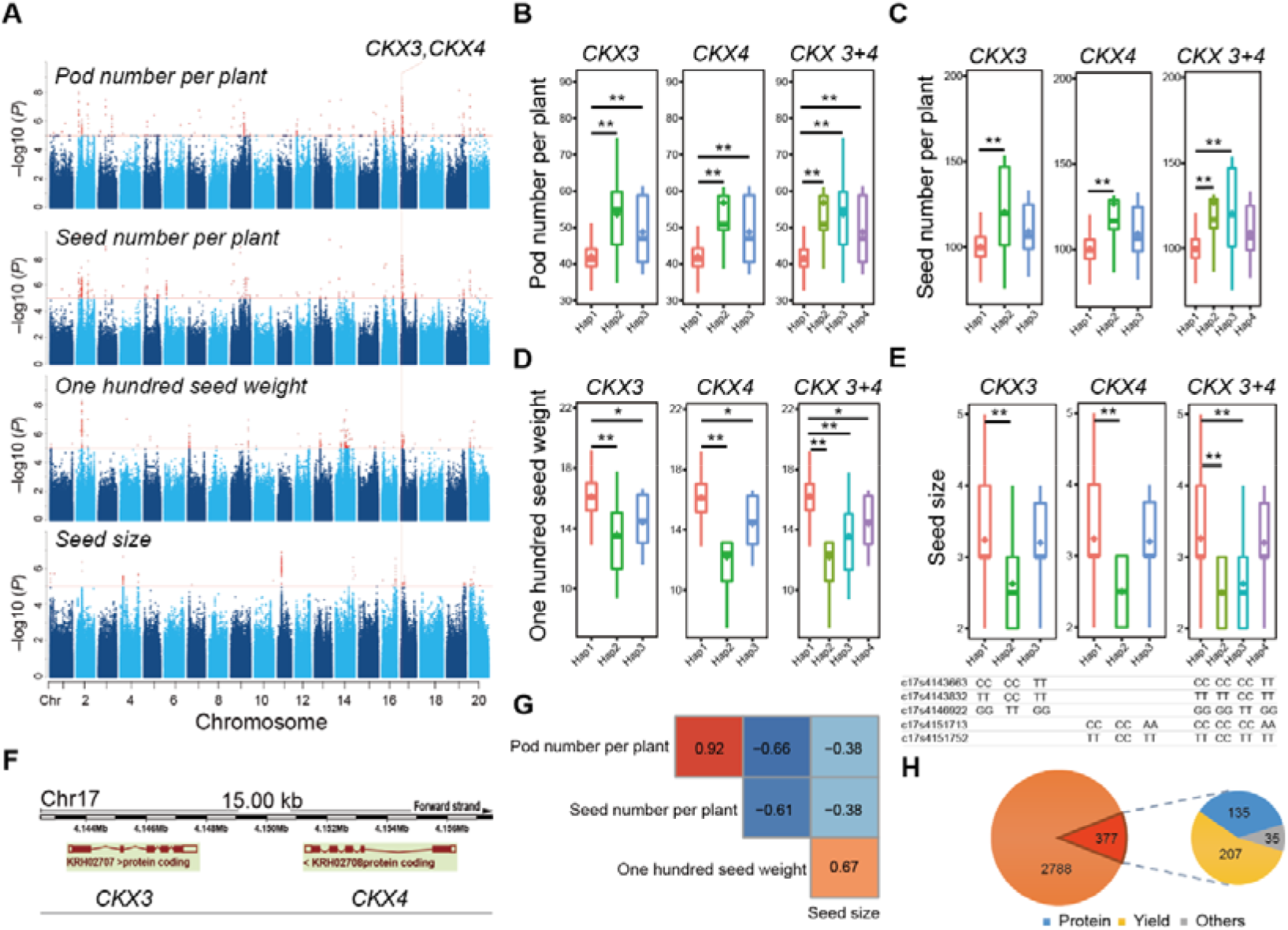
Association of *CKX* and yield related traits in soybean. **A.** Manhantan plot of four yield related traits pod number per plant, seed number per plant, one hundred seed weight and seed size, and the candidate *CKX* genes. **B.** Pod number per plant distribution for the haplotypes of *CKX* genes.**C.** Seed number per plant distribution for the haplotypes of *CKX* genes. **D.** One hundred seed weight distribution for the haplotypes of *CKX* genes. **E.** Seed size distribution for the haplotypes of *CKX* genes. **F.** Chromosome location and transcripts structure of *CKX3* and *CKX4*. **G.** Phenotype correlation of four traits. **H.** Statistics of SAGs located in selective sweep regions and their percentage for trait categories. (\**P* < 0.05; \*\**P* < 0.01)

We further analysed the relationship between functional mutations in *CKX3* and *CKX4*. There were three non-synonymous SNPs on *CKX3* and two non-synonymous SNPs on *CKX4*. As these two genes are only approximately 3 kb apart from each other in the same genome region, we analysed the two genes separately as well as combined as one in relation to their association with haplotypes and traits (**Figure 4B-E**). Results showed that the functional mutations can both form three haplotypes for *CKX3* and *CKX4 separately*, and four haplotypes for *CKX3+4* combined. For all comparisons in all traits, *Haplotype 1* always showed significant differences compared with the other haplotypes. The relationship between different haplotypes in terms of pod or seed number per plant showed a consistent trend, while that of one hundred seed weight or seed size showed a consistent but opposite trend. There was a phenotypic correlation between pod number per plant and seed number per plant (0.92), and between one hundred seed weight and seed size (0.67) (**Figure 4F**). Furthermore, we observed that *CKX3* and *CKX4* were located in different strands of the same chromosome, suggesting they are more likely to have independent functions. However, we did not detect expression of *CKX4* in the subsequent qPCR validation. Thus, only *CKX3* is regarded as a real candidate gene, while the role of *CKX4* need further study.

When we compared the soybean accession information of each haplotype for the four yield-related traits, we observed that most accessions with the *Haplotype 1* genotype had dominant traits (lower pod or seed numbers and larger seeds and seed weights), and were more associated with cultivars. The other haplotypes were mainly landrace-specific haplotypes, and these accessions all belong to *Group 1*. We found that *CKX3* was also located on a strong selective sweep region. This indicates that the functional mutation sites in *CKX3* experienced strong directed artificial selection, resulting in genotype differences and affecting yield related traits. Furthermore, we compared all SAGs and selective sweep regions for all traits, and found that approximately 12% of genes are located in the selected sweep regions, which have experienced artificial selection (**Supplementary Table 9**). It is interesting to note that of all the SAGs located in selective sweep regions, about 55% are related to yield traits, 36% related to protein traits, and less than 10% are related to other traits (**Figure 4G, H**).

### CYP85A2 is associated with architecture and yield traits

There is one SAL on chromosome 18 which is associated with six traits including plant height, main stem number, stem strength, lodging, podding habit, and seed weight per plant. Interestingly, these traits include both architecture and yield related traits. A cytochrome P450 family gene named *CYP85A2* is located within a 4.37 kb region of this SAL (**Figure 5A, B**). The association of the *CPY85A2* gene with architecture and yield traits in soybean is a novel finding. We also observed that a non-synonymous mutation site *c18s55526062* is involved in producing two haplotypes. The haplotype with a *CC* genotype has a dwarf plant height, a low main stem node number, a high stem strength and a low lodging rate. When plants produced mostly limited or semi-limited pods, their seed weight per plant was also found to increase (**Figure 5C**). Phenotypically, plant height was positively correlated with main stem node number (0.95), while stem strength and lodging were negatively correlated (−0.81), showing a trend consistent with the genotype (**Figure 5D**). The *CC* genotype of *c18s55526062* is a dominant genotype, which is useful when designing an ideal plant type and increasing soybean yield.

**Figure 5.**
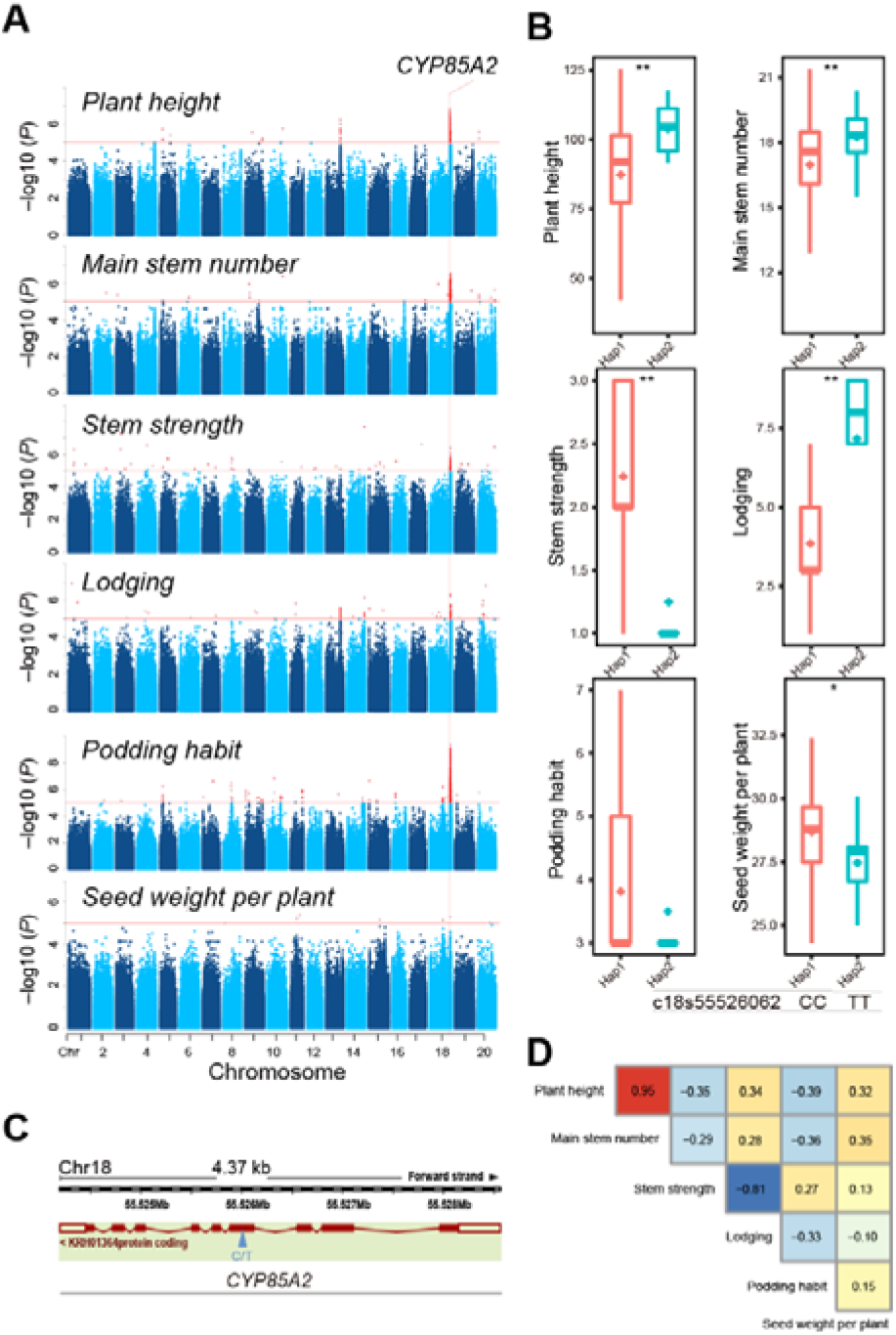
Association of *CYP85A2* and architecture or yield related traits in soybean. **A.** Manhantan plot of plant height, main stem number, stem strength, lodging, podding habit, seed weight per plant, and the candidate gene *CYP85A2*. **B.** Traits distribution for the haplotypes of gene *CYP85A2*. **C.** Chromosome location and transcripts structure of *CYP85A2.* **D.** Phenotype correlation of the six traits. (\**P* < 0.05; \*\**P* < 0.01)

### Different phenotypes coupled through hub gene modules form a complex phenotype-gene network

Based on in-depth exploration of the GWAS results, we observed that one trait is associated with multiple genes and vice versa. At the same time, due to widespread protein-level interactions between genes, a complex network was also found between various phenotypes and genes. In order to explore this further, we used a functional mutation-based haplotype test to screen SAGs in all SALs for all traits. We then constructed a phenotype-gene network which included 34 traits and 853 SAGs (**Figure 6**). At the trait level, they were divided into six categories, namely architecture, colour, oil, isoflavone, protein, and yield. At the gene level, besides the six categories, there emerged a mixed category with which genes associated more than with any one trait category. We found that traits in the same category were closely linked within the entire network. However, some trait categories were also linked with each other, such as yield, oil, protein and colour, and they were all closely linked to architecture through common SAGs. This suggests that there are subtle relationships between architecture and other trait categories. In this genetic network, six trait categories were linked through 15 hub nodes containing a total of 367 genes (**Supplementary Table 10**). The largest hub was *HA1* (short form for *Hub Architecture 1*). The genes in this hub were only associated with two or more architecture-related traits. Unlike *HA1*, the genes in the *HA2* node were associated with two or more architecture-related traits, but also had protein interactions with other genes. Multiple yield traits were associated with hub *HY1*, containing *CKX3*, while hub *HM4*, which contains *CYP85A2*, was connected with architecture and yield traits.

**Figure 6.**
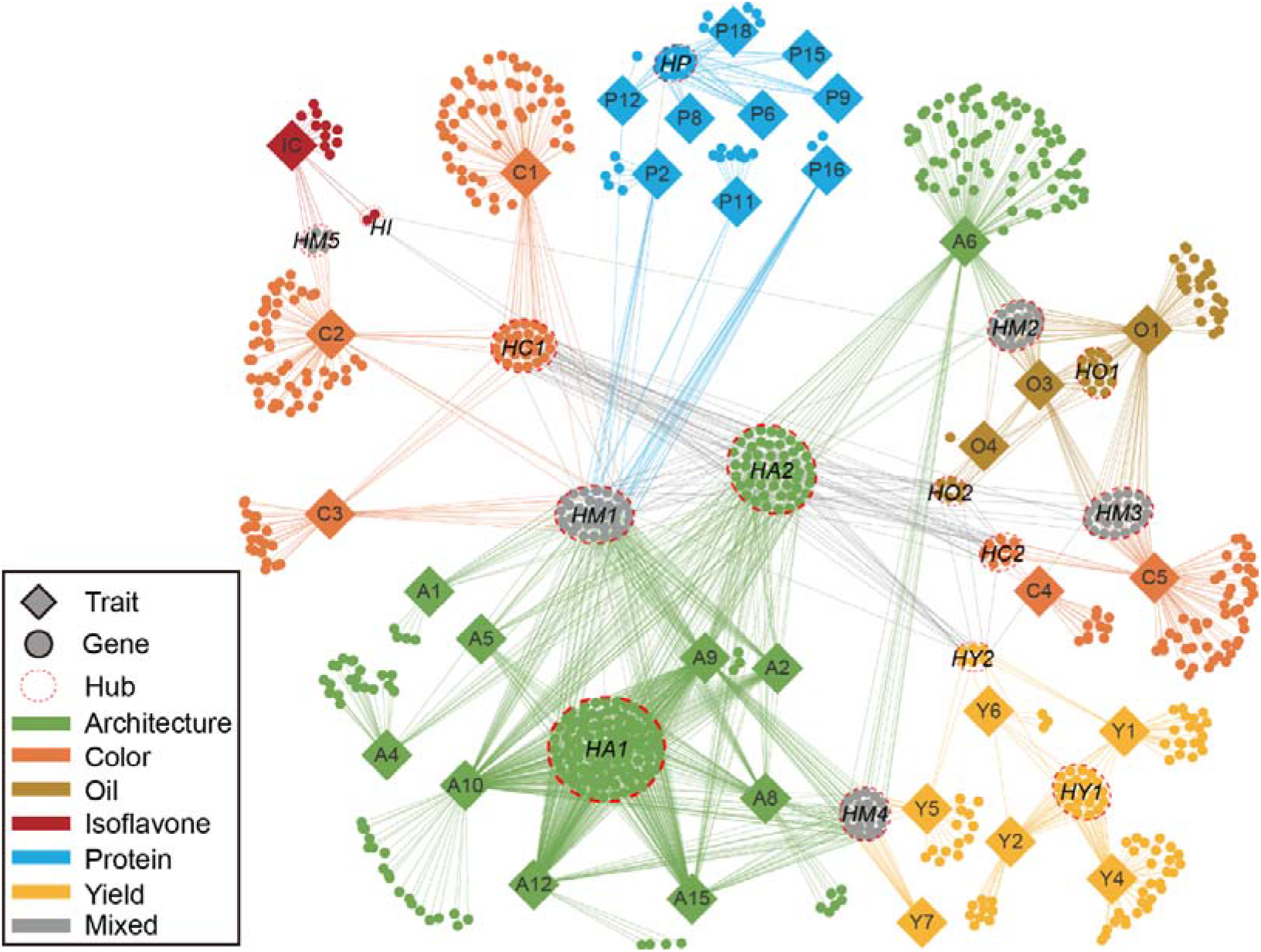
Phenotype-gene genetic network in soybean. Traits are solid rhombuses, genes are solid circles, and hubs are hollow ellipses. Six trait categories, their associated genes and links between them are colored accordingly, genes associated with more than one trait categories are colored grey. Genes with protein-protein interaction are linked with gray lines.

## Discussion

In this study, we deeply sequenced 250 representative landrace and cultivar soybean accessions. Through population genetics and GWAS analyses, the genetic structure of European soybean varieties was analysed for the first time. Novel candidate genes related to seed isoflavone content, yield and architecture traits were identified. Moreover, we constructed a soybean phenotype-gene interaction network, and found evidence of the improvement of soybean yield related traits at molecular level.

A total of ~3T bases and 6M SNPs were obtained, the maximum sequencing depth of a single accession was 22.5x, with the average depth at 11x (higher than previous soybean resequencing studies[2–4]. Eighty-four percent of the accessions with their genome were sequenced for the first time, which provides new data for soybean genome research. Previous soybean research mainly focused on varieties from Asia and North America, but not Europe[3, 4]. This study completed the resequencing of 26 European accessions and, for the first time, outlines a breeding history of European soybean. It was found that European soybean cultivars had higher genetic diversities and lower breeding levels compared to North American cultivars. Both European and American soybean cultivars may have been introduced from different ancestors in China. This theory is based on the following findings: there is a small population difference between European varieties and Chinese landraces, and between American varieties and Chinese cultivars, whilst there is a large population difference between American and European varieties (**Figure 2G, Supplementary Figure 2**). Our findings are consistent with the current hypothesis that soybean originated in China, and they show that ancestral components from the area of origin are the most complex. This study showed that the heterozygosity rates of most accessions are less than 0.2, except four accessions with a higher heterozygosity which may be caused by their complex ancestral compositions (**Supplementary Table S3**). Further combination analysis of the selective sweep and GWAS revealed that artificial selection of soybean at the phenotypic level is consistent with the genome level. Genomic regions associated with yield and quality traits are more likely to experience artificial selection. This may be a reflection of yield- and quality-directed artificial selection of soybean breeding at the genetic level. Furthermore, evidence of functional mutations under artificial selection for a candidate gene *CKX3* related to multiple yield traits were identified. The results of this study provide valuable information for marker-assisted selection, which is vital in the improvement of soybean breeding.

Isoflavone is a secondary metabolite produced via phenylpropane metabolic pathways in higher plants. Isoflavone is associated with plant stress resistance, defence against microbial and insect infection, promotion of rhizobium chemotaxis, and the development of rhizome and nitrogen fixation in plants. It also provides health benefits to human, such as in reducing the incidence of cancer and cardiovascular diseases, and regulating the immune response[12]. Therefore, increasing the seed isoflavone content of soybean can improve its nutritional and health benefits. However, few genome-wide studies have investigated the molecular mechanism of soybean’s isoflavone content. Isoflavone is synthesized in the cytoplasm, but due to cell cytotoxicity, it cannot accumulate in the cytoplasm and must be continuously transported to vacuoles for storage. Therefore, isoflavone content mainly depends on two factors: synthesis efficiency and transport efficiency[13]. The transcript factor *GL3* is a *bHLH* gene family member which can form the MYB-bHLH-WD40 (MBW) complex with two other transcription factors (*MYB* and *WD40*) to jointly regulate the synthesis of flavonoids and anthocyanin in plants[14]. *GST* can bind with glutathione (GSH) to form an ABC transporter to transport and catalyse the entry of flavonoids into vacuoles for accumulation[13]. In this study, we identified four novel genes that may be associated with isoflavone content. These genes include transcription factors *GL3*, which participate in the regulation of multi-enzyme systems from phenylpropanoid to isoflavone biosynthesis pathways, and two *GST* genes, *GSTT1* and *GSTL3*, which facilitate the transporting of isoflavone from the cytoplasm to vacuoles (**Figure 3F**). In addition, we observed many other genes in the SALs, such as cation/H+ exchanger (*CHX20*), pyrophosphorylase 4 (*PPa4*), an actin depolymerizing factor 7 (*ADF7*), a mitochondrial substrate carrier family protein, a myosin heavy chain-related protein, an *ATP* synthase alpha/beta family protein, and a protein kinase superfamily protein, among others (**Supplementary Table 8**) are all related to isoflavone transport. This over-representation of transport-related genes further suggests that the accumulation of soybean isoflavone is related to its transport to the vacuole. In conclusion, soybean isoflavone content is not merely determined by one or several genes or loci, but by a multiple gene system involved in synthesis, regulation, transport, and storage.

We observed that other novel candidate genes, such as *CKX3,* is associated with multiple yield traits. We also observed, for the first time, that *CYP85A2* is associated with multiple architecture and yield traits in soybean. It is well-known that cytokinin promotes cell division and plant growth, and *CKX* is one of the key enzymes in cytokinin metabolism. A functional variation in the *CKX* gene may affect the cytokinin metabolism, thus affecting grain yield and related traits. A number of studies on *Arabidopsis thaliana*, rice and other crops have shown that mutation or reduced expression levels of *CKX* family genes are related to a decrease in seed setting rate and an increase in seed weight[15, 16]. *CYP85A2* is involved in the brassinosteroid biosynthesis pathway in *Arabidopsis thaliana* and it converts 6-deoxocastasterone to castasterone, which is followed by the conversion of castasterone to brassinolide[17]. Brassinosteroids (BRs) are broad-spectrum plant growth regulators, playing an important role in plant growth and development, as well as in biological and abiotic stress responses[18]. Mutations in the *CYP85A2* gene have led to an increased production of the dwarf phenotype[19], and an overexpression of the *CYP85A* family gene resulting in increased BR content, biomass, plant height, plant fresh weight and fruit yield[20]. These results showed that, *CKX3* and *CYP85A2* may affect soybean yield and architecture related traits through different molecular mechanisms. The potential effect of functional mutations in these genes on the phenotypes was further confirmed by our haplotype tests. However, to verify whether these candidate genes and functional mutations are the true cause of the phenotypic differences, further functional verification of these genes is necessary. Multiple methods such as the construction of isolated populations, transgene, gene knockout, gene editing, and expression verification could be used for this purpose. In this study, we performed expression verification in seedlings with different haplotypes/phenotypes for six genes *GL3*, *GSTL3*, *GSTT1b*, *CKX3*, *CKX4* and *CYP85A2.* The results show that, except for no expression were detected for *CKX4*, all the other five genes were expressed differently for different haplotypes/phenotypes in seedlings; the expression levels of *GL3*, *GSTL3* and *GSTT1b* related to isoflavone content in the strains with high isoflavone content values was significantly higher than that in the strains with low isoflavone content values (T-test, P<0.05); the expression level of *CKX3* in the strains with high yield phenotype values was significantly higher than that in the strains with low yield phenotype values (T-test, P<0.05) (**Supplementary Figure 55**).

The highest goal of plant breeding is to aggregate many desired traits into a single genome. Breeders need to simultaneously select and improve multiple related traits. However, because multiple traits are interrelated, it is possible that when screening for a favourable trait one also selects an unfavourable one. Understanding the genetic network behind different traits can help breeders increase breeding efficiency. Although soybean genetic networks for multiple agronomic traits have been established at the loci level[4], we built a new phenotype-gene network which includes 34 traits and 853 genes. This network reflects the relationship between phenotypes and genes more directly than the previous phenotype-SAL network, and is more conducive to the discovery of important candidate genes. For example, the *Hub Mixed 1* (*HM1*) node was associated with two or more trait types (architecture, colour or protein), while the *HBT* gene in the *HM1* node was associated with six architectural traits (branch number, main stem number, plant height, stem strength, lodging, and podding habit) and four protein-related traits (phenylalanine content, isoleucine content, tyrosine content, and glycine content). It is known that the *HBT* gene belongs to the *CDC27b* gene family and is involved in cell cycle regulation, which is related to cell development and division[21]. Therefore, soybean architecture is likely affected by *HBT*, despite its relationship with amino acid content is unclear. There are also many other interesting examples of the above. Leaf shape is known to affect photosynthesis efficiency, followed by carbohydrate accumulation, and, as a consequence, oil accumulation, while Hub *HM2*, containing *FAD2*, connects oil content and leaf shape[22]. It has also been reported that oil traits and seed coat lustre traits experienced parallel selection during bean domestication[23]. The hub *HM3* node connects oil-related traits and seed coat lustre. Anthocyanin synthesis and isoflavone synthesis share part of their metabolic pathways and hub *HM5* connects colour traits and isoflavone content. Our phenotype-gene network may surpass the phenotype-SAL network in terms of candidate gene selection, which is also beneficial to polymerization breeding programs. For example, breeders can achieve polymerization breeding by directly selecting a favourable gene (such as *CYP85A2*) in Hub *HM4*, which is related to both yield and architecture traits, and eliminate the confusion of other adverse genes located in the same SAL. Furthermore, it is worth noting that the architecture related traits, which centrally connect various other trait categories, have the most extensive connectivity. In other words, there are numerous relationships between architecture related traits and other trait categories in the phenotype-gene network (**Figure 6**), suggesting that some candidate genes related to architecture traits may also be related to other trait types. This may provide theoretical support and practical guidance for parallel selection breeding and promote “ideotype” breeding in soybean. The next step is to conduct more in-depth functional investigations on genes with a potential application value, such as *CKX3* and *CYP85A2*. This would help promote the design and breeding process of soybean varieties with a higher yield and quality. Overall, our work is conducive to promoting soybean genome functional research and genomic breeding.

## Materials and Methods

### Plant materials and phenotyping

A total of 250 soybean varieties were analysed in this study, which were provided by the National Crop Germplasm Resources Platform, Institute of Crop Genetics, Institute of Crop Sciences, Chinese Academy of Agricultural Sciences. All materials were planted and phenotyped at three locations: the Gongzhuling experimental site in the Jilin Academy of Agricultural Sciences (latitude 43.51°, east longitude 124.80°), the Harbin experimental site in the Heilongjiang Academy of Agricultural Sciences (45.68° north latitude, 126.61° east longitude), and the Chifeng experimental site un the Agricultural Science Institute in Inner Mongolia (42.27° north latitude, 118.90° east longitude) in late April of 2008, 2009 and 2010, respectively. Grain protein content was measured using the Kjeldahl method from National Food Safety Standard GB5009.5-2010 China[24], while the grain fatty acid contents were determined using the Soxhlet extraction method from National Food Safety Standard GB/T5512-2008, China[25]. Amino acid content was determined using high performance liquid chromatography (S433D, Seckam, Germany) following a previous amino acid determination method from National Food Safety Standard GB/T 18246-2000, China[26]. Grain isoflavone content was determined using high performance liquid chromatography from National Food Safety Standard GB/T23788-2009, China[27]. Finally, the phenotypic data were integrated using the BLUP (Best Linear Unbiased Prediction) method using R[28] in order to remove environmental effects and obtain stable genetic phenotypes. Seeds were planted in (CLC-BIV-M/CLC404-TV, MMM, Germany) at 20°C (with 12-h day/12-h night) and a relative humidity of 60–80% till six leaves stage (about two-week-old). Two-week-old seedlings (24°C, 12-h day/12-h night cycle) were used in this qPCR validation.

### DNA preparation and sequencing

The genomic DNA for all soybean accessions were extracted from soybean leaves after three weeks of growth. DNA extraction was performed using the cetyltrimethylammonium bromide (CTAB) method[29]. The library for each accession was constructed with an insert size of approximately 500 base pairs, following manufacturer’s instructions (Illumina Inc., San Diego, CA, USA). All soybean accessions were sequenced and paired-end 150 bp reads were produced using an Illumina NovaSeq 6000 sequencer at the BerryGenomics Company (http://www.berrygenomics.com/ Beijing, China).

### Total RNA extraction, cDNA synthesis and qRT-PCR analysis

Total RNA was isolated for each sample using TRIzol Reagent (Invitrogen, Nottingham, UK) according to the manufacturer’s instructions. The purified RNA was stored at −80°C until subsequent analyses. According to the manufacturer’s instructions (Takara, Shiga, Japan), first-strand cDNA synthesis was performed using M-MLV reverse transcriptase. Quantitative real-time PCR (qRT-PCR) was performed using a SYBR Premix Ex Taq Kit (Takara) and a real-time PCR machine (CFX96; Bio-Rad, Hercules, CA, USA), following the manufacturer’s instructions. The procedure used for qRT-PCR was 95°C at 10 minutes, followed by 38 cycles of 15 s at 95°C and 60 s at 61-62°C. β-actin was used as a reference gene for analysis of relative expression patterns of mRNA. The reactions were carried out with three biological replicates, with at least two technical replicates for each sample. The data were analyzed using the method according to the previous study[30], and the means ± standard errors (SE) of three biological replicates are presented.

### Mapping, variant calling and annotation

Raw paired-end resequencing reads were first cleaned by removing reads with adaptors, reads of low quality and reads with “N”s. The high-quality clean reads were then mapped to the soybean reference genome (Williams 82 assembly v2.1) with BWA[31]. Statistical analyses of mapping rate and genomic coverage of clean reads were performed using in-house scripts. The Speedseq pipeline[32] was used for SNP and Indel calling, and vcftools[33] was used for genotype filtering. Missing genotypes were imputed and phased through a localized haplotype clustering algorithm implemented using Beagle v3.0[34]. Variant annotation was performed using ANNOVAR[35] against the soybean gene model set v2.1.42. After annotation, SNPs and Indels were categorized into exonic, intronic, intergenic, splicing, 5′UTRs, 3′UTRs, upstream, and downstream. Exonic SNPs were further categorized into synonymous, nonsynonymous, stop gain, and stop loss. Exonic Indels were further categorized into frameshift, non-frameshift, stop gain, and stop loss.

### Population structure analysis

Approximately 6M SNPs from the 250 soybean accessions were concatenated for the construction of a phylogenetic tree. Using a neighbour joining algorithm with a pairwise gap deletion method for 100 bootstrap replications, a phylogenetic tree was constructed with MegaCC[36]. The output was displayed using the iTOL[37] web tool. With the whole genome genotype as the input, a principal component analysis (PCA) was done using flashPCA [38] and the first two eigenvectors were plotted. A population admixture analysis with *k = 2* to *k = 5* parameters were set to infer the admixture of ancestors using fastSTRUCTURE.

### Genetic diversity analysis

Linkage disequilibrium analyses for each subpopulation were performed using PLINK[39] by calculating the correlation coefficient (*r^2^*) of any two SNP pairs in one chromosome. An LD decay plot was drawn using the average *r^2^* value for the distance from 0 to 1,000 kb. Pairwise IBS calculations were also performed using PLINK and a distance matrix was generated for each subpopulation. Population genetic diversities were measured using VCFtools[33] by calculating *θπ* and *Fst*. *θπ* was used to measure the genetic diversity of each subpopulation, while *Fst*, plus the allele frequency distribution (AFD) plot (which was generated by in-house scripts), were used to measure genetic diversity between subpopulations. In addition, sliding window calculations of *r^2^, θπ*, *Fst* and *Tajima’ D* values were also performed for genome-wide displays of soybean genetic diversities with a 100 kb window and a 10 kb step.

### Selective sweep analysis

We used two methods to detect selective sweep regions across the soybean genome: *Tajima’ D* combined *θπ* and *Fst* combined *θπ* ratios. Firstly, a genome-wide sliding window calculation of *θπ*, *Fst*, and *Tajima’ D* values (with a 100 kb window and a 10 kb step) were performed on landraces, cultivars, and the whole population, respectively. Secondly, the top 5% of the *Tajima’ D* and *θπ* windows for the whole population were selected. In addition, the top 5% of the *Fst* and *θπ* ratio windows for the landraces versus cultivars were also selected. Thirdly, the selected windows from these two methods were merged together to become the final selective sweep regions. ROH analyses for each accession were performed using PLINK[39] with the parameters of a minimum ROH length set to 300 kb.

### GWAS and significantly associated loci

Association analysis for each trait on each SNP with an MAF larger than 0.05 was performed using a single-locus mixed linear model (MLM) implemented in GEMMA[40] (which corrects confounding by population structure and the relatedness matrix). The GWAS results were displayed using a Manhattan plot and a QQ-plot created with the R package CMplot[41]. A clump based method implemented in PLINK[39] was used to reduce a false peak and to detect real SALs. The P-value cut-off was set to 10^−5^ so as to, firstly, uncover significant associated SNPs. Following this, for each significantly associated SNP, if there were more than 10 SNPs within a 100 kb distance that had P-values smaller than 10^−4^, then the region was regarded as a potential SAL. Finally, all overlapping SALs were merged to generate final SAL sets and the SNP with the smallest P-value in a SAL was defined as a peak.

### Detection of significantly associated genes

There are usually tens of genes in a SAL, and it is difficult to determine which genes are truly associated with traits and which are irrelevant. We improved a functional mutation-based haplotype test method for SAG discovery in SAL. As most variants within a gene are non-functional, the gene’s amino acid sequence and its function will not change. Only a few variants have the potential to change a gene’s amino acid sequence, such as nonsynonymous SNPs, frameshift Indels, variants in splicing sites, promoter regions, start codons, and stop codons. These combined functional mutations can only produce two or three different gene haplotypes. It is possible to test the relationship between gene haplotypes and traits. If they are significantly associated, then the gene is also most likely associated with the trait, which is how SAG is defined. In this study, Welch’s test was used for a two-group haplotype test and a Tukey’s test was used for a multiple group haplotype test to detect SAGs. Functional annotation of SAGs was directly retrieved from SoyBase[42].

### Network construction

Of all the genes located in the SALs, the most significant SAGs with a P-value smaller than 10^−5^, and their corresponding traits, were retained to build the phenotype-gene network for soybean. Protein-protein interaction information for soybean was retrieved form the String database[43] and mapped to the soybean genes using BLAST[44]. Construction, visualization and exploration of the network was performed using Cytoscape[45].

**Table 1.** Functional variants of representative significant associated genes

| Variant ID | Chrom | Positon | Ref | Alt | Variant type | Gene ID | Gene<br>symbol |
| --- | --- | --- | --- | --- | --- | --- | --- |
| c5s38936266 | 5 | 38936266 | C | T | nonsynonymous SNV | GLYMA_05G206900 | *GSTT1a* |
| c5s38940717 | 5 | 38940717 | C | T | nonsynonymous SNV | GLYMA_05G207000 | *GSTT1b* |
| c5s39035509 | 5 | 39035509 | G | C | nonsynonymous SNV | GLYMA_05G208300 | *GL3* |
| c5s39036346 | 5 | 39036346 | T | C | nonsynonymous SNV | GLYMA_05G208300 | *GL3* |
| c13s24804891 | 13 | 24804891 | C | T | nonsynonymous SNV | GLYMA_13G135600 | *GSTL3* |
| c13s24805363 | 13 | 24805363 | A | T | splicing SNV | GLYMA_13G135600 | *GSTL3* |
| c17s4143663 | 17 | 4143663 | C | T | nonsynonymous SNV | GLYMA_17G054500 | *CKX3* |
| c17s4143832 | 17 | 4143832 | T | C | nonsynonymous SNV | GLYMA_17G054500 | *CKX3* |
| c17s4146922 | 17 | 4146922 | G | T | nonsynonymous SNV | GLYMA_17G054500 | *CKX3* |
| c17s4151713 | 17 | 4151713 | C | A | nonsynonymous SNV | GLYMA_17G054600 | *CKX4* |
| c17s4151752 | 17 | 4151752 | T | C | nonsynonymous SNV | GLYMA_17G054600 | *CKX4* |
| c18s55526062 | 18 | 55526062 | C | T | nonsynonymous SNV | GLYMA_18G272300 | *CYP85A2* |

## Data availability

The raw sequence data reported in this paper have been deposited in the Genome Sequence Archive[46] in BIG Data Center[47], Beijing Institute of Genomics (BIG), Chinese Academy of Sciences, under accession numbers CRA002552 that are publicly accessible at https://bigd.big.ac.cn/gsa. The variation data reported in this paper have been deposited in the Genome Variation Map [48] under accession number GVM000076 that can be publicly accessible at http://bigd.big.ac.cn/gvm/getProjectDetail?project=GVM000076. The bioinformatics analysis scripts used in this paper can be download through https://github.com/yjthu/GPB_250SoyReseq.

## CRediT statement

**Chunming Yang:** Resources, Investigation, Validation. **Jun Yan:** Methodology, Formal analysis, Writing, Revision. **Shuqin Jiang:** Formal analysis. **Xia Li:** Investigation, Validation. **Haowei Min:** Conceptualization, Supervision, Formal analysis, Writing, Revision. **Xiangfeng Wang:** Conceptualization, Supervision, Writing, Revision. **Dongyun Hao:** Conceptualization, Supervision, Writing, Revision. All authors read and approved the final manuscript.

## Acknowledgements

This research was supported by grants from the Agricultural Science and Technology Innovation Project, Jilin Province [CXGC2017ZY027], and from Program of Accurate Identification and Display of Soybean Germplasm, China [NB08-2130315-(25-31)-06, NB07-2130315-(25-30)-06, NB06-070401-(22-27)-05, NB2010-2130315-25-05]. We are grateful to Prof. Lijuan Qiu for her agreement of using the 250 soybean varieties from her laboratory at China Academy of Agricultural Sciences. We appreciate Dr. Zhangxiong Liu of China Academy of Agricultural Sciences for the technical guidance in soybean phenotypic characterization. Acknowledgement also goes to Yunshan Wei(Inner Mongolia Academy of Agriculture & Animal Husbandry Sciences), Shuhong Wei and Qiang Wang(Heilongjiang Academy of Agricultural Sciences), for partially phenotypic characterization of the soybean population used in this work.

## Supplementary material

Supplementary Tables 1-10; Supplementary Figures 1-55.

**Supplementary Table 1** Summary of mapping and coverage

**Supplementary Table 2** Summary of SNPs and Indels

**Supplementary Table 3** The ancestry proportion estimates for each accession

**Supplementary Table 4** Putative regions experiencing selective sweeps

**Supplementary Table 5** Summary of ROH regions in soybean varieties

**Supplementary Table 6** Information of 50 agronomic traits

**Supplementary Table 7** Summary of significant associated loci detected by GWAS analysis

**Supplementary Table 8** Genes located in significant associated loci for isoflavone content

**Supplementary Table 9** Genes located in both significant associated loci and putative selective sweep regions

**Supplementary Table 10** Summary of hub genes in soybean agronomic traits networks

**Figure S1** SNP density distribution across soybean chromosomes.

**Figure S2** Allele frequency distribution between soybean subpopulations.

**Figure S3** Selective sweep analysis for 250 soybean accessions. **A.** Selective sweep analysis by *Tajima’D* combine *θπ*. **B.** Selective sweep analysis by *Fst* combine *θπ* ratios. Red dots present the top 5% selected windows. **C.** Venn diagram of genes screened by two selective sweep analysis methods. **D.** Venn diagram of genes screened by two selective sweep analysis methods and ROH analysis.

**Figure S4** Phenotype correlations between 50 soybean traits.

**Figure S5** GWAS of pod height at bottom using MLM. A. Density distribution of pod height at bottom. B. Manhattan plots for pod height at bottom. Negative log10 P-values from a genome-wide scan are plotted against SNP positions of 20 chromosomes. C. Quantile-quantile plot for pod height at bottom. The horizontal red line indicates the significant threshold (10-5). Trait-associated SNPs above the significant threshold are colored in red.

**Figure S6** GWAS of effective branch number using MLM. A. Density distribution of effective branch number. B. Manhattan plots for effective branch number. Negative log10 P-values from a genome-wide scan are plotted against SNP positions of 20 chromosomes. C. Quantile-quantile plot for effective branch number. The horizontal red line indicates the significant threshold (10-5). Trait-associated SNPs above the significant threshold are colored in red.

**Figure S7** GWAS of pubescence density using MLM. **A.** Density distribution of pubescence density. **B.** Manhattan plots for pubescence density. Negative log_10_ P-values from a genome-wide scan are plotted against SNP positions of 20 chromosomes. **C.** Quantile-quantile plot for pubescence density. The horizontal red line indicates the significant threshold (10^−5^). Trait-associated SNPs above the significant threshold are colored in red.

**Figure S8** GWAS of defollation using MLM. **A.** Density distribution of defollation. **B.** Manhattan plots for defollation. Negative log_10_ P-values from a genome-wide scan are plotted against SNP positions of 20 chromosomes. **C.** Quantile-quantile plot for defollation. The horizontal red line indicates the significant threshold (10^−5^). Trait-associated SNPs above the significant threshold are colored in red.

**Figure S9** GWAS of inflorenscence length using MLM. **A.** Density distribution of inflorenscence length. **B.** Manhattan plots for inflorenscence length. Negative log_10_ P-values from a genome-wide scan are plotted against SNP positions of 20 chromosomes. **C.** Quantile-quantile plot for inflorenscence length. The horizontal red line indicates the significant threshold (10^−5^). Trait-associated SNPs above the significant threshold are colored in red.

**Figure S10** GWAS of leaf shape using MLM. **A.** Density distribution of leaf shape. **B.** Manhattan plots for leaf shape. Negative log_10_ P-values from a genome-wide scan are plotted against SNP positions of 20 chromosomes. **C.** Quantile-quantile plot for leaf shape. The horizontal red line indicates the significant threshold (10^−5^). Trait-associated SNPs above the significant threshold are colored in red.

**Figure S11** GWAS of leaflet size using MLM. **A.** Density distribution of leaflet size. Manhattan plots for leaflet size. Negative log_10_ P-values from a genome-wide scan are plotted against SNP positions of 20 chromosomes. **C.** Quantile-quantile plot for leaflet size. The horizontal red line indicates the significant threshold (10^−5^). Trait-associated SNPs above the significant threshold are colored in red.

**Figure S12** GWAS of lodging using MLM. **A.** Density distribution of lodging. **B.** Manhattan plots for lodging. Negative log_10_ P-values from a genome-wide scan are plotted against SNP positions of 20 chromosomes. **C.** Quantile-quantile plot for lodging. The horizontal red line indicates the significant threshold (10^−5^). Trait-associated SNPs above the significant threshold are colored in red.

**Figure S13** GWAS of number of nodes on main stem using MLM. **A.** Density distribution of number of nodes on main stem. **B.** Manhattan plots for number of nodes on main stem. Negative log_10_ P-values from a genome-wide scan are plotted against SNP positions of 20 chromosomes. **C.** Quantile-quantile plot for number of nodes on main stem. The horizontal red line indicates the significant threshold (10^−5^). Trait-associated SNPs above the significant threshold are colored in red.

**Figure S14** GWAS of plant height using MLM. **A.** Density distribution of plant height. **B.** Manhattan plots for plant height. Negative log_10_ P-values from a genome-wide scan are plotted against SNP positions of 20 chromosomes. **C.** Quantile-quantile plot for plant height. The horizontal red line indicates the significant threshold (10^−5^). Trait-associated SNPs above the significant threshold are colored in red.

**Figure S15** GWAS of plant type using MLM. **A.** Density distribution of plant type. **B.** Manhattan plots for plant type. Negative log_10_ P-values from a genome-wide scan are plotted against SNP positions of 20 chromosomes. **C.** Quantile-quantile plot for plant type. The horizontal red line indicates the significant threshold (10^−5^). Trait-associated SNPs above the significant threshold are colored in red.

**Figure S16** GWAS of stem termination using MLM. **A.** Density distribution of podding habit. **B.** Manhattan plots for stem termination. Negative log_10_ P-values from a genome-wide scan are plotted against SNP positions of 20 chromosomes. **C.** Quantile-quantile plot for stem termination. The horizontal red line indicates the significant threshold (10^−5^). Trait-associated SNPs above the significant threshold are colored in red.

**Figure S17** GWAS of seed crack using MLM. **A.** Density distribution of seed crack. **B.** Manhattan plots for seed crack. Negative log_10_ P-values from a genome-wide scan are plotted against SNP positions of 20 chromosomes. **C.** Quantile-quantile plot for seed crack. The horizontal red line indicates the significant threshold (10^−5^). Trait-associated SNPs above the significant threshold are colored in red.

**Figure S18** GWAS of stem diameter using MLM. **A.** Density distribution of stem diameter. **B.** Manhattan plots for stem diameter. Negative log_10_ P-values from a genome-wide scan are plotted against SNP positions of 20 chromosomes. **C.** Quantile-quantile plot for stem diameter. The horizontal red line indicates the significant threshold (10^−5^). Trait-associated SNPs above the significant threshold are colored in red.

**Figure S19** GWAS of stem intension using MLM. **A.** Density distribution of stem intension. **B.** Manhattan plots for stem intension. Negative log_10_ P-values from a genome-wide scan are plotted against SNP positions of 20 chromosomes. **C.** Quantile-quantile plot for stem intension. The horizontal red line indicates the significant threshold (10^−5^). Trait-associated SNPs above the significant threshold are colored in red.

**Figure S20** GWAS of pubescence color using MLM. **A.** Density distribution of pubescence color. **B.** Manhattan plots for pubescence color. Negative log_10_ P-values from a genome-wide scan are plotted against SNP positions of 20 chromosomes. **C.** Quantile-quantile plot for pubescence color. The horizontal red line indicates the significant threshold (10^−5^). Trait-associated SNPs above the significant threshold are colored in red.

**Figure S21** GWAS of flower color using MLM. **A.** Density distribution of flower color. **B.** Manhattan plots for flower color. Negative log_10_ P-values from a genome-wide scan are plotted against SNP positions of 20 chromosomes. **C.** Quantile-quantile plot for flower color. The horizontal red line indicates the significant threshold (10^−5^). Trait-associated SNPs above the significant threshold are colored in red.

**Figure S22** GWAS of leaf color using MLM. **A.** Density distribution of leaf color. **B.** Manhattan plots for leaf color. Negative log_10_ P-values from a genome-wide scan are plotted against SNP positions of 20 chromosomes. **C.** Quantile-quantile plot for leaf color. The horizontal red line indicates the significant threshold (10^−5^). Trait-associated SNPs above the significant threshold are colored in red.

**Figure S23** GWAS of mature pod color using MLM. **A.** Density distribution of mature pod color. **B.** Manhattan plots for mature pod color. Negative log_10_ P-values from a genome-wide scan are plotted against SNP positions of 20 chromosomes. **C.** Quantile-quantile plot for mature pod color. The horizontal red line indicates the significant threshold (10^−5^). Trait-associated SNPs above the significant threshold are colored in red.

**Figure S24** GWAS of seed coat luster using MLM. **A.** Density distribution of seed coat luster. **B.** Manhattan plots for seed coat luster. Negative log_10_ P-values from a genome-wide scan are plotted against SNP positions of 20 chromosomes. **C.** Quantile-quantile plot for seed coat luster. The horizontal red line indicates the significant threshold (10^−5^). Trait-associated SNPs above the significant threshold are colored in red.

**Figure S25** GWAS of isoflavone content using MLM. **A.** Density distribution of isoflavone content. **B.** Manhattan plots for isoflavone content. Negative log_10_ P-values from a genome-wide scan are plotted against SNP positions of 20 chromosomes. **C.** Quantile-quantile plot for isoflavone content. The horizontal red line indicates the significant threshold (10^−5^). Trait-associated SNPs above the significant threshold are colored in red.

**Figure S26** GWAS of linoleic acid content using MLM. **A.** Density distribution of linoleic acid content. **B.** Manhattan plots for linoleic acid content. Negative log_10_ P-values from a genome-wide scan are plotted against SNP positions of 20 chromosomes. **C.** Quantile-quantile plot for linoleic acid content. The horizontal red line indicates the significant threshold (10^−5^). Trait-associated SNPs above the significant threshold are colored in red.

**Figure S27** GWAS of linolenic acid content using MLM. **A.** Density distribution of linolenic acid content. **B.** Manhattan plots for linolenic acid content. Negative log_10_ P-values from a genome-wide scan are plotted against SNP positions of 20 chromosomes. **C.** Quantile-quantile plot for linolenic acid content. The horizontal red line indicates the significant threshold (10^−5^). Trait-associated SNPs above the significant threshold are colored in red.

**Figure S28** GWAS of oleic acid content using MLM. **A.** Density distribution of oleic acid content. **B.** Manhattan plots for oleic acid content. Negative log_10_ P-values from a genome-wide scan are plotted against SNP positions of 20 chromosomes. **C.** Quantile-quantile plot for oleic acid content. The horizontal red line indicates the significant threshold (10^−5^). Trait-associated SNPs above the significant threshold are colored in red.

**Figure S29** GWAS of palmitic acid content using MLM. **A.** Density distribution of palmitic acid content. **B.** Manhattan plots for palmitic acid content. Negative log_10_ P-values from a genome-wide scan are plotted against SNP positions of 20 chromosomes. **C.** Quantile-quantile plot for palmitic acid content. The horizontal red line indicates the significant threshold (10^−5^). Trait-associated SNPs above the significant threshold are colored in red.

**Figure S30** GWAS of crude protein content using MLM. **A.** Density distribution of crude protein content. **B.** Manhattan plots for crude protein content. Negative log_10_ P-values from a genome-wide scan are plotted against SNP positions of 20 chromosomes. **C.** Quantile-quantile plot for crude protein content. The horizontal red line indicates the significant threshold (10^−5^). Trait-associated SNPs above the significant threshold are colored in red.

**Figure S31** GWAS of alanine content using MLM. **A.** Density distribution of alanine content. **B.** Manhattan plots for alanine content. Negative log_10_ P-values from a genome-wide scan are plotted against SNP positions of 20 chromosomes. **C.** Quantile-quantile plot for alanine content. The horizontal red line indicates the significant threshold (10^−5^). Trait-associated SNPs above the significant threshold are colored in red.

**Figure S32** GWAS of arginine content using MLM. **A.** Density distribution of arginine content. **B.** Manhattan plots for arginine content. Negative log_10_ P-values from a genome-wide scan are plotted against SNP positions of 20 chromosomes. **C.** Quantile-quantile plot for arginine content. The horizontal red line indicates the significant threshold (10^−5^). Trait-associated SNPs above the significant threshold are colored in red.

**Figure S33** GWAS of aspartic acid content using MLM. **A.** Density distribution of aspartic acid content. **B.** Manhattan plots for aspartic acid content. Negative log_10_ P-values from a genome-wide scan are plotted against SNP positions of 20 chromosomes. **C.** Quantile-quantile plot for aspartic acid content. The horizontal red line indicates the significant threshold (10^−5^). Trait-associated SNPs above the significant threshold are colored in red.

**Figure S34** GWAS of glutamate content using MLM. **A.** Density distribution of glutamate content. **B.** Manhattan plots for glutamate content. Negative log_10_ P-values from a genome-wide scan are plotted against SNP positions of 20 chromosomes. **C.** Quantile-quantile plot for glutamate content. The horizontal red line indicates the significant threshold (10^−5^). Trait-associated SNPs above the significant threshold are colored in red.

**Figure S35** GWAS of glycine content using MLM. **A.** Density distribution of glycine content. **B.** Manhattan plots for glycine content. Negative log_10_ P-values from a genome-wide scan are plotted against SNP positions of 20 chromosomes. **C.** Quantile-quantile plot for glycine content. The horizontal red line indicates the significant threshold (10^−5^). Trait-associated SNPs above the significant threshold are colored in red.

**Figure S36** GWAS of histidine content using MLM. **A.** Density distribution of histidine content. **B.** Manhattan plots for histidine content. Negative log_10_ P-values from a genome-wide scan are plotted against SNP positions of 20 chromosomes. **C.** Quantile-quantile plot for histidine content. The horizontal red line indicates the significant threshold (10^−5^). Trait-associated SNPs above the significant threshold are colored in red.

**Figure S37** GWAS of isoleucine content using MLM. **A.** Density distribution of isoleucine content. **B.** Manhattan plots for isoleucine content. Negative log_10_ P-values from a genome-wide scan are plotted against SNP positions of 20 chromosomes. **C.** Quantile-quantile plot for isoleucine content. The horizontal red line indicates the significant threshold (10^−5^). Trait-associated SNPs above the significant threshold are colored in red.

**Figure S38** GWAS of leucine content using MLM. **A.** Density distribution of leucine content. **B.** Manhattan plots for leucine content. Negative log_10_ P-values from a genome-wide scan are plotted against SNP positions of 20 chromosomes. **C.** Quantile-quantile plot for leucine content. The horizontal red line indicates the significant threshold (10^−5^). Trait-associated SNPs above the significant threshold are colored in red.

**Figure S39** GWAS of lysine content using MLM. **A.** Density distribution of lysine content. **B.** Manhattan plots for lysine content. Negative log_10_ P-values from a genome-wide scan are plotted against SNP positions of 20 chromosomes. **C.** Quantile-quantile plot for lysine content. The horizontal red line indicates the significant threshold (10^−5^). Trait-associated SNPs above the significant threshold are colored in red.

**Figure S40** GWAS of methionine content using MLM. **A.** Density distribution of methionine content. **B.** Manhattan plots for methionine content. Negative log_10_ P-values from a genome-wide scan are plotted against SNP positions of 20 chromosomes. **C.** Quantile-quantile plot for methionine content. The horizontal red line indicates the significant threshold (10^−5^). Trait-associated SNPs above the significant threshold are colored in red.

**Figure S41** GWAS of phenylalanine content using MLM. **A.** Density distribution of phenylalanine content. **B.** Manhattan plots for phenylalanine content. Negative log_10_ P-values from a genome-wide scan are plotted against SNP positions of 20 chromosomes. **C.** Quantile-quantile plot for phenylalanine content. The horizontal red line indicates the significant threshold (10^−5^). Trait-associated SNPs above the significant threshold are colored in red.

**Figure S42** GWAS of proline content using MLM. **A.** Density distribution of proline content. **B.** Manhattan plots for proline content. Negative log_10_ P-values from a genome-wide scan are plotted against SNP positions of 20 chromosomes. **C.** Quantile-quantile plot for proline content. The horizontal red line indicates the significant threshold (10^−5^). Trait-associated SNPs above the significant threshold are colored in red.

**Figure S43** GWAS of serine content using MLM. **A.** Density distribution of serine content. **B.** Manhattan plots for serine content. Negative log_10_ P-values from a genome-wide scan are plotted against SNP positions of 20 chromosomes. **C.** Quantile-quantile plot for serine content. The horizontal red line indicates the significant threshold (10^−5^). Trait-associated SNPs above the significant threshold are colored in red.

**Figure S44** GWAS of threonine content using MLM. **A.** Density distribution of threonine content. **B.** Manhattan plots for threonine content. Negative log_10_ P-values from a genome-wide scan are plotted against SNP positions of 20 chromosomes. **C.** Quantile-quantile plot for threonine content. The horizontal red line indicates the significant threshold (10^−5^). Trait-associated SNPs above the significant threshold are colored in red.

**Figure S45** GWAS of tyrosine content using MLM. **A.** Density distribution of tyrosine content. **B.** Manhattan plots for tyrosine content. Negative log_10_ P-values from a genome-wide scan are plotted against SNP positions of 20 chromosomes. **C.** Quantile-quantile plot for tyrosine content. The horizontal red line indicates the significant threshold (10^−5^). Trait-associated SNPs above the significant threshold are colored in red.

**Figure S46** GWAS of valine content using MLM. **A.** Density distribution of valine content. **B.** Manhattan plots for valine content. Negative log_10_ P-values from a genome-wide scan are plotted against SNP positions of 20 chromosomes. **C.** Quantile-quantile plot for valine content. The horizontal red line indicates the significant threshold (10^−5^). Trait-associated SNPs above the significant threshold are colored in red.

**Figure S47** GWAS of total amino acids content using MLM. **A.** Density distribution of total amino acids content. **B.** Manhattan plots for total amino acids content. Negative log_10_ P-values from a genome-wide scan are plotted against SNP positions of 20 chromosomes. **C.** Quantile-quantile plot for total amino acids content. The horizontal red line indicates the significant threshold (10^−5^). Trait-associated SNPs above the significant threshold are colored in red.

**Figure S48** GWAS of hundred grain weight using MLM. **A.** Density distribution of one hundred seed weight. **B.** Manhattan plots for hundred grain weight. Negative log_10_ P-values from a genome-wide scan are plotted against SNP positions of 20 chromosomes. **C.** Quantile-quantile plot for hundred grain weight. The horizontal red line indicates the significant threshold (10^−5^). Trait-associated SNPs above the significant threshold are colored in red.

**Figure S49** GWAS of pod number per plant using MLM. **A.** Density distribution of pod number per plant. **B.** Manhattan plots for pod number per plant. Negative log_10_ P-values from a genome-wide scan are plotted against SNP positions of 20 chromosomes. **C.** Quantile-quantile plot for pod number per plant. The horizontal red line indicates the significant threshold (10^−5^). Trait-associated SNPs above the significant threshold are colored in red.

**Figure S50** GWAS of pod size using MLM. **A.** Density distribution of pod size. **B.** Manhattan plots for pod size. Negative log_10_ P-values from a genome-wide scan are plotted against SNP positions of 20 chromosomes. **C.** Quantile-quantile plot for pod size. The horizontal red line indicates the significant threshold (10^−5^). Trait-associated SNPs above the significant threshold are colored in red.

**Figure S51** GWAS of seed number per plant using MLM. **A.** Density distribution of seed number per plant. **B.** Manhattan plots for seed number per plant. Negative log_10_ P-values from a genome-wide scan are plotted against SNP positions of 20 chromosomes. **C.** Quantile-quantile plot for seed number per plant. The horizontal red line indicates the significant threshold (10^−5^). Trait-associated SNPs above the significant threshold are colored in red.

**Figure S52** GWAS of seed number per pod using MLM. **A.** Density distribution of seed number per pod. **B.** Manhattan plots for seed number per pod. Negative log_10_ P-values from a genome-wide scan are plotted against SNP positions of 20 chromosomes. **C.** Quantile-quantile plot for seed number per pod. The horizontal red line indicates the significant threshold (10^−5^). Trait-associated SNPs above the significant threshold are colored in red.

**Figure S53** GWAS of seed size using MLM. **A.** Density distribution of seed size. **B.** Manhattan plots for seed size. Negative log_10_ P-values from a genome-wide scan are plotted against SNP positions of 20 chromosomes. **C.** Quantile-quantile plot for seed size. The horizontal red line indicates the significant threshold (10^−5^). Trait-associated SNPs above the significant threshold are colored in red.

**Figure S54** GWAS of seed weight per plant using MLM. **A.** Density distribution of seed weight per plant. **B.** Manhattan plots for seed weight per plant. Negative log_10_ P-values from a genome-wide scan are plotted against SNP positions of 20 chromosomes. **C.** Quantile-quantile plot for seed weight per plant. The horizontal red line indicates the significant threshold (10^−5^). Trait-associated SNPs above the significant threshold are colored in red.

**Figure S55** Gene expression validation of different haplotypes/phenotypes for five candidate genes. **A.** Phenotype distribution (left) and expression level (right) of *GL3* for different haplotypes. **B.** Phenotype distribution (left) and expression level (right) of *GSTL3* for different haplotypes. **C.** Phenotype distribution (left) and expression level (right) of *GSTT1b* for different haplotypes. **D.** Phenotype distribution (left) and expression level (right) of *CKX3* for different haplotypes. **E.** Phenotype distribution (left) and expression level (right) of *CYP85A2* for different haplotypes.

