## Supplemental Figures for "Resequencing 250 soybean accessions: new insights into genes associated with agronomic traits and genetic networks"

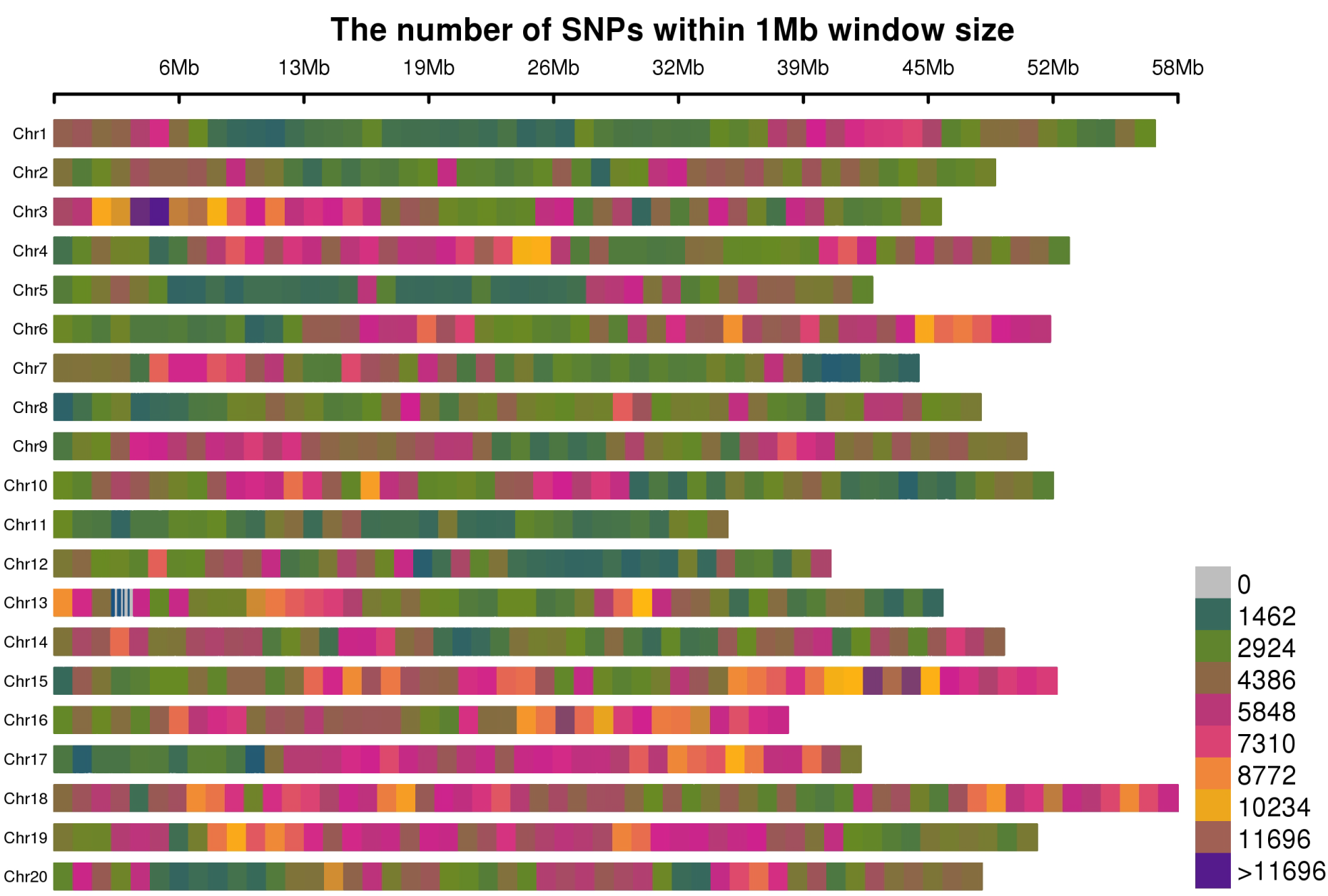

**Figure S1** SNP density distribution across soybean chromosomes.

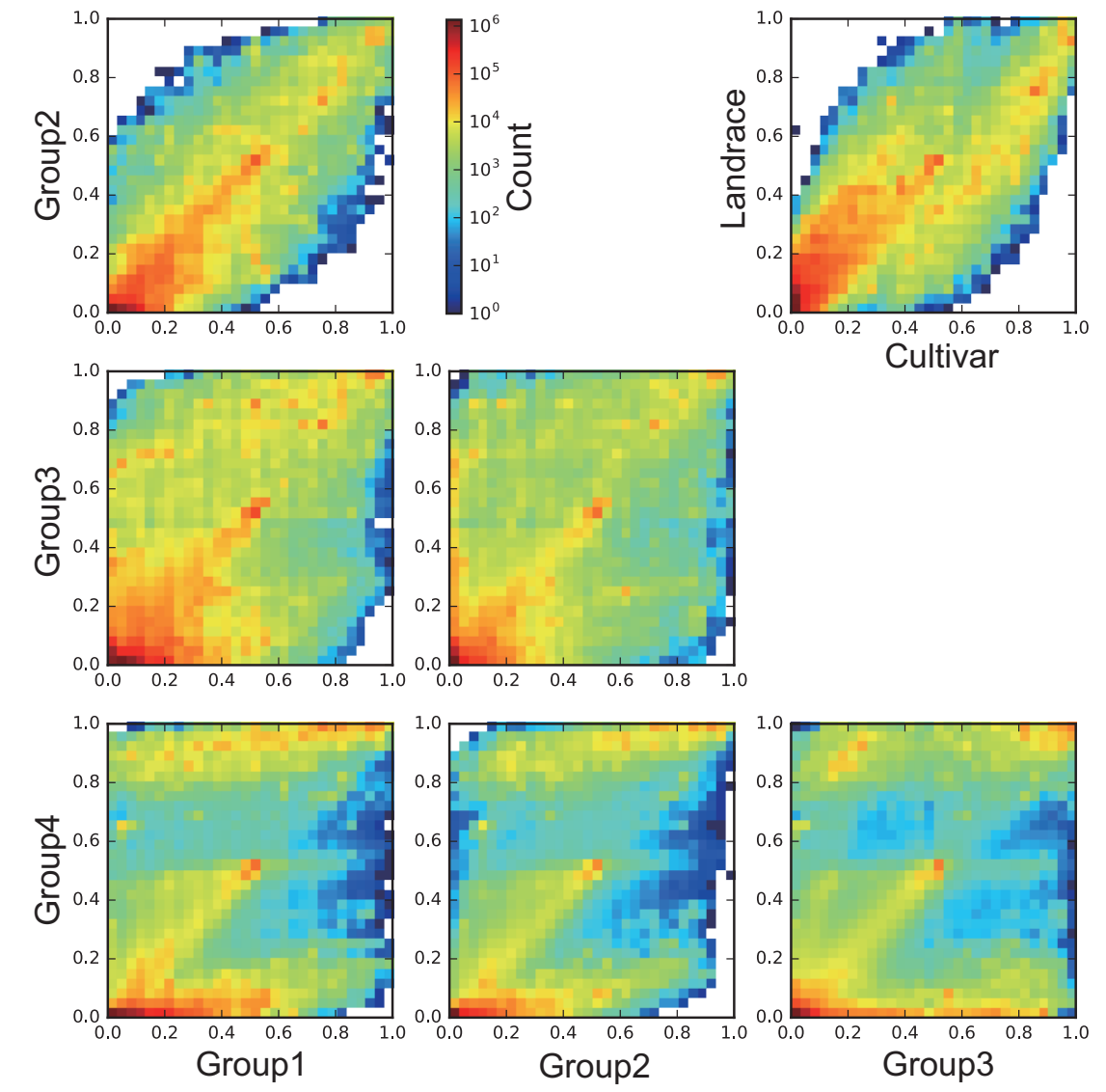

**Figure S2** Allele frequency distribution between soybean subpopulations.

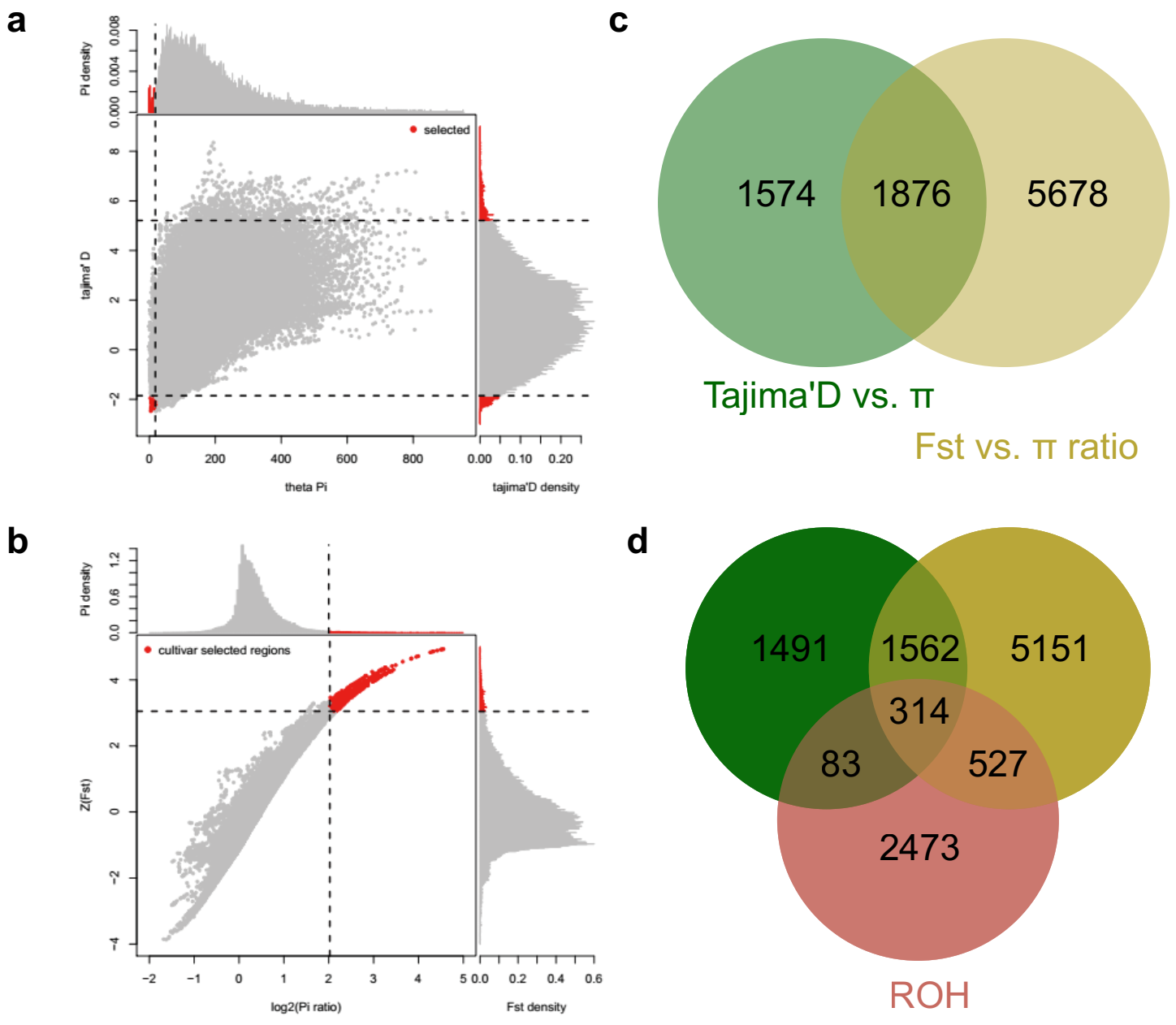

**Figure S3** Selective sweep analysis for 250 soybean accessions. **A.** Selective sweep analysis by *Tajima’D* combine *θπ.* **B.** Selective sweep analysis by *Fst* combine *θπ* ratios. Red dots present the top 5% selected windows. **C.** Venn diagram of genes screened by two selective sweep analysis methods. **D.** Venn diagram of genes screened by two selective sweep analysis methods and ROH analysis.

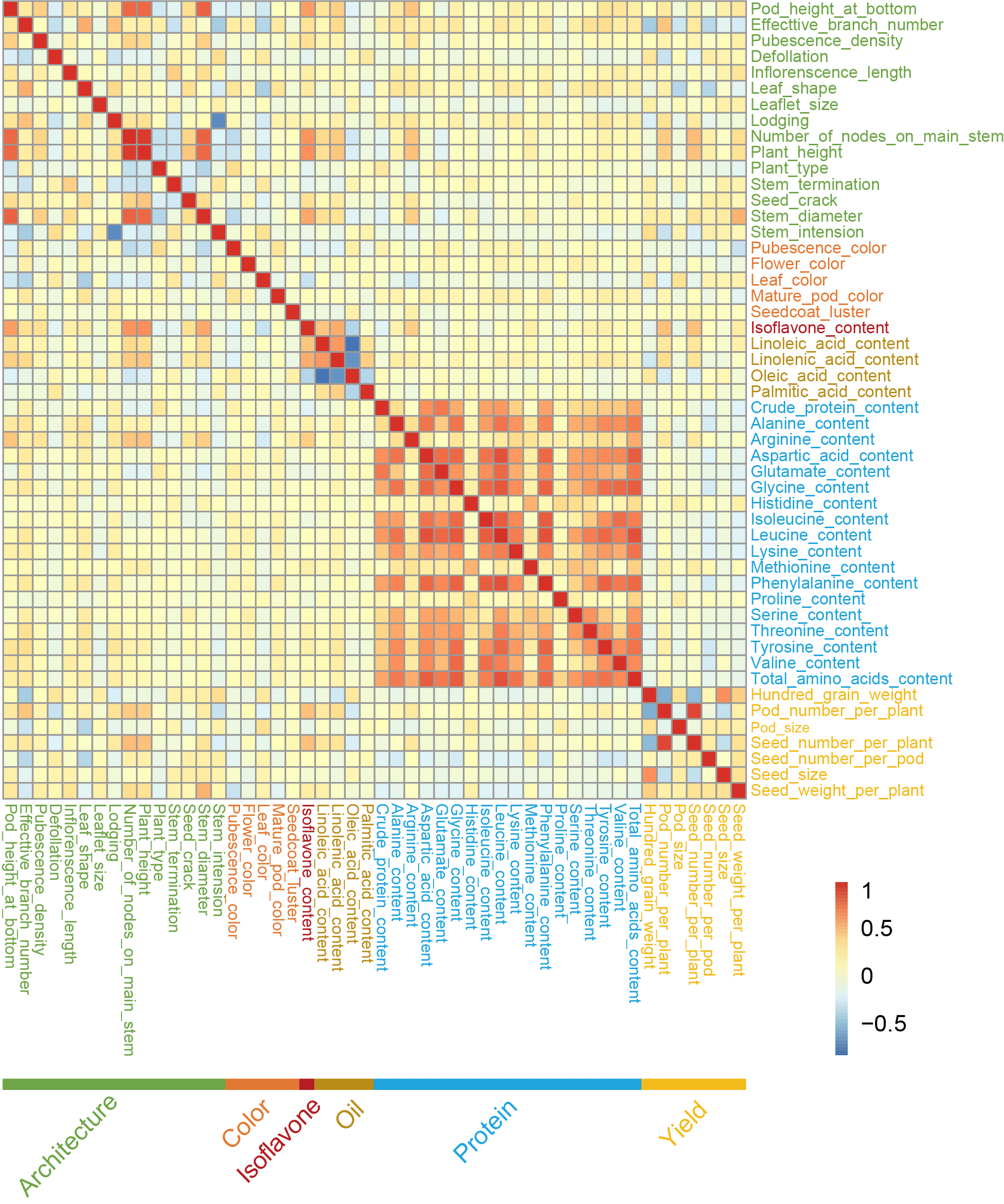

**Figure S4** Phenotype correlations between 50 soybean traits.

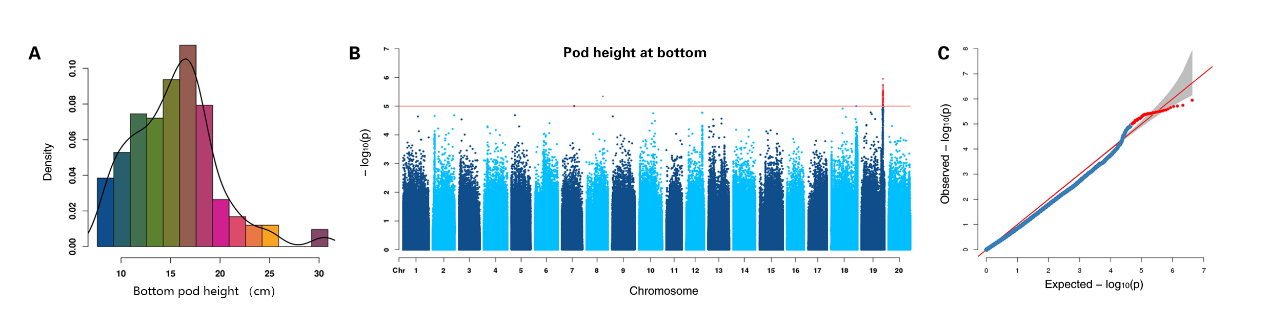

**Figure S5** GWAS of pod height at bottom using MLM. **A.** Density distribution of pod height at bottom. **B.** Manhattan plots for pod height at bottom. Negative log_10_ P-values from a genome-wide scan are plotted against SNP positions of 20 chromosomes. **C.** Quantile-quantile plot for pod height at bottom. The horizontal red line indicates the significant threshold (10^-5^). Trait-associated SNPs above the significant threshold are colored in red.

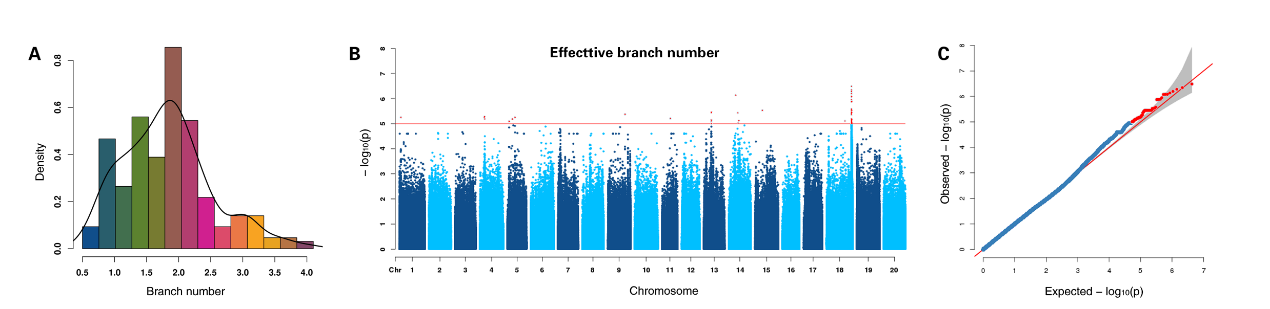

**Figure S6** GWAS of effective branch number using MLM. **A.** Density distribution of effective branch number. **B.** Manhattan plots for effective branch number. Negative log_10_ P-values from a genome-wide scan are plotted against SNP positions of 20 chromosomes. **C.** Quantile-quantile plot for effective branch number. The horizontal red line indicates the significant threshold (10^-5^). Trait-associated SNPs above the significant threshold are colored in red.

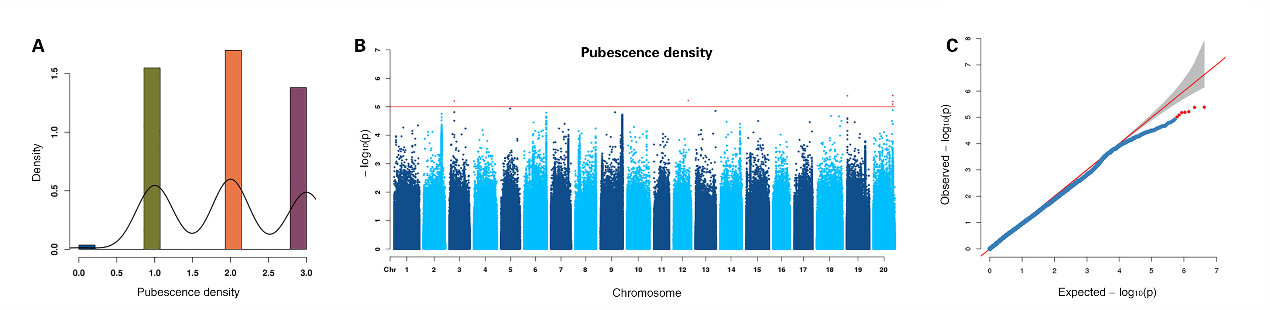

**Figure S7** GWAS of pubescence density using MLM. **A.** Density distribution of pubescence density. **B.** Manhattan plots for pubescence density. Negative log_10_ P-values from a genome-wide scan are plotted against SNP positions of 20 chromosomes. **C.** Quantile-quantile plot for pubescence density. The horizontal red line indicates the significant threshold (10^-5^). Trait-associated SNPs above the significant threshold are colored in red.

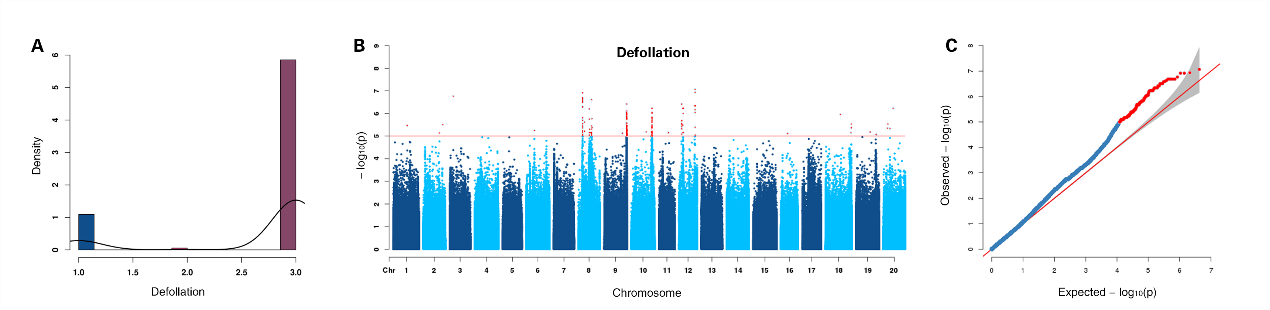

**Figure S8** GWAS of defollation using MLM. **A.** Density distribution of defollation. **B.** Manhattan plots for defollation. Negative log_10_ P-values from a genome-wide scan are plotted against SNP positions of 20 chromosomes. **C.** Quantile-quantile plot for defollation. The horizontal red line indicates the significant threshold (10^-5^). Trait-associated SNPs above the significant threshold are colored in red.

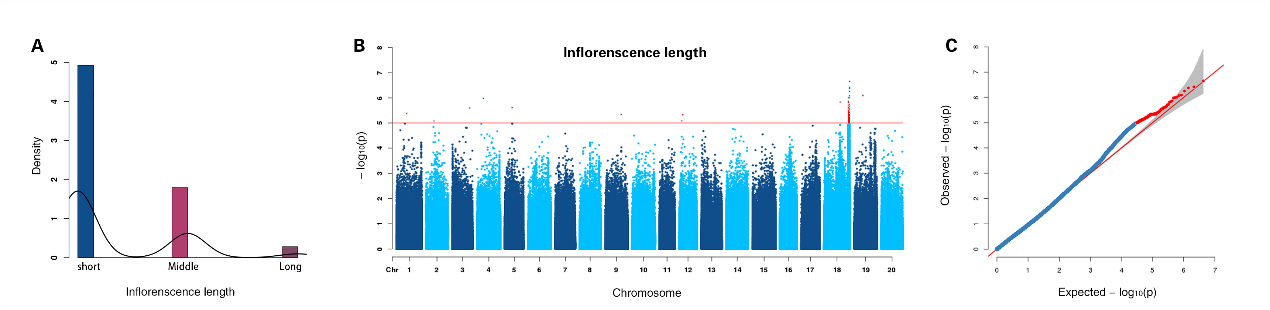

**Figure S9** GWAS of inflorenscence length using MLM. **A.** Density distribution of inflorenscence length. **B.** Manhattan plots for inflorenscence length. Negative log_10_ P-values from a genome-wide scan are plotted against SNP positions of 20 chromosomes. **C.** Quantile-quantile plot for inflorenscence length. The horizontal red line indicates the significant threshold (10^-5^). Trait-associated SNPs above the significant threshold are colored in red.

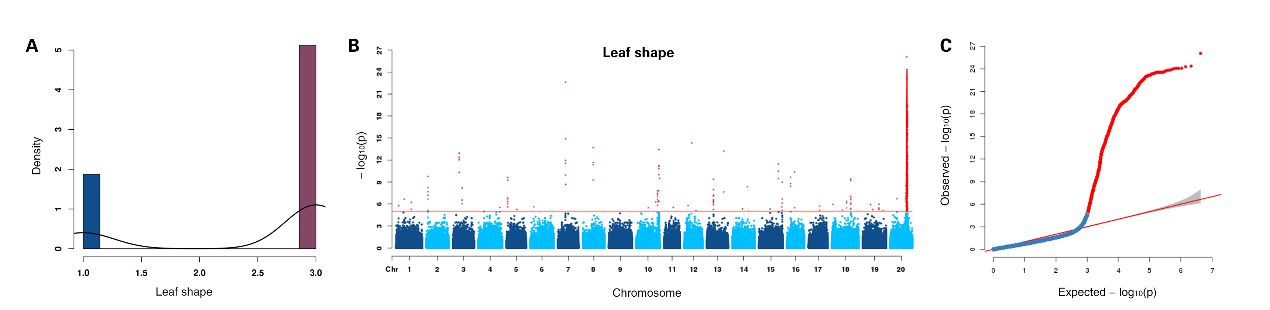

**Figure S10** GWAS of leaf shape using MLM. **A.** Density distribution of leaf shape. **B.** Manhattan plots for leaf shape. Negative log_10_ P-values from a genome-wide scan are plotted against SNP positions of 20 chromosomes. **C.** Quantile-quantile plot for leaf shape. The horizontal red line indicates the significant threshold (10^-5^). Trait-associated SNPs above the significant threshold are colored in red.

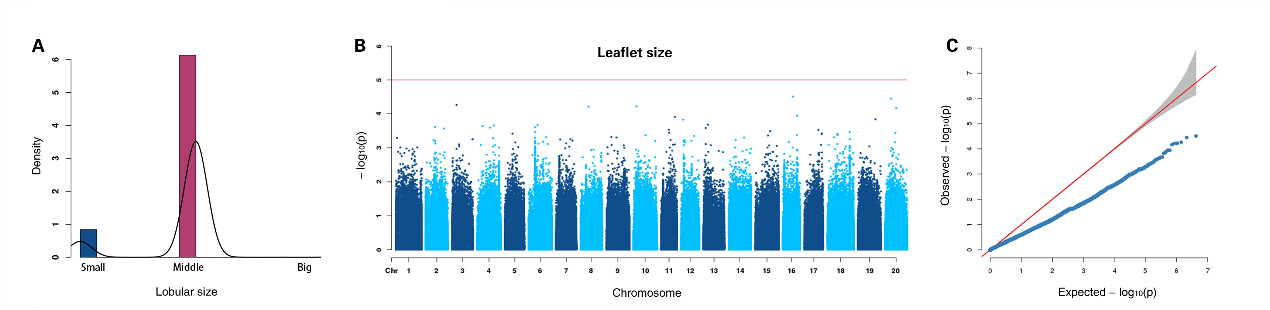

**Figure S11** GWAS of leaflet size using MLM. **A.** Density distribution of leaflet size. **B.** Manhattan plots for leaflet size. Negative log_10_ P-values from a genome-wide scan are plotted against SNP positions of 20 chromosomes. **C.** Quantile-quantile plot for leaflet size. The horizontal red line indicates the significant threshold (10^-5^). Trait-associated SNPs above the significant threshold are colored in red.

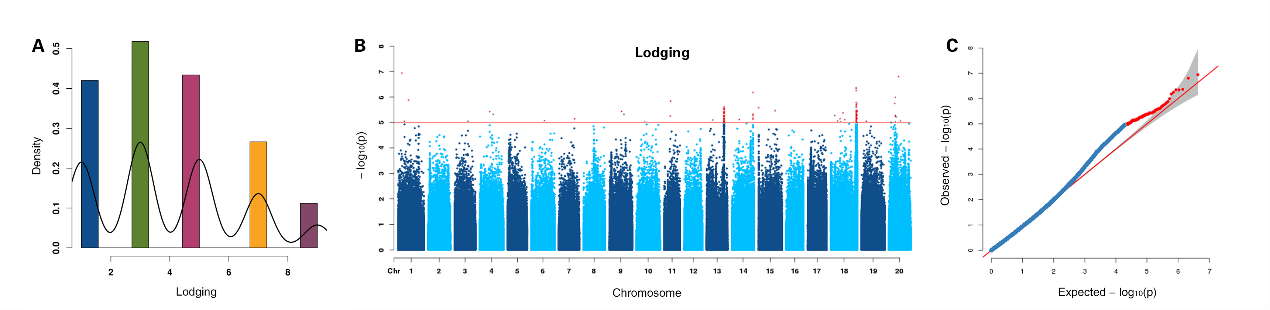

**Figure S12** GWAS of lodging using MLM. **A.** Density distribution of lodging. **B.** Manhattan plots for lodging. Negative log_10_ P-values from a genome-wide scan are plotted against SNP positions of 20 chromosomes. **C.** Quantile-quantile plot for lodging. The horizontal red line indicates the significant threshold (10^-5^). Trait-associated SNPs above the significant threshold are colored in red.

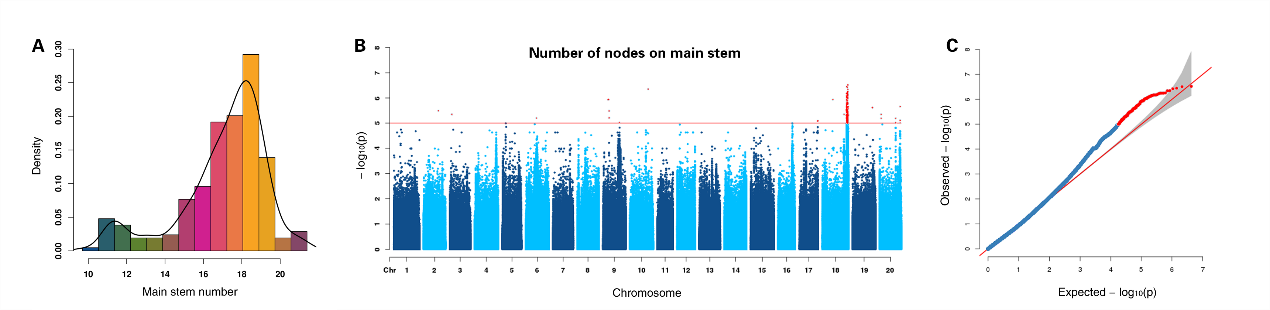

**Figure S13** GWAS of number of nodes on main stem using MLM. **A.** Density distribution of number of nodes on main stem. **B.** Manhattan plots for number of nodes on main stem. Negative log_10_ P-values from a genome-wide scan are plotted against SNP positions of 20 chromosomes. **C.** Quantile-quantile plot for number of nodes on main stem. The horizontal red line indicates the significant threshold (10^-5^). Trait-associated SNPs above the significant threshold are colored in red.

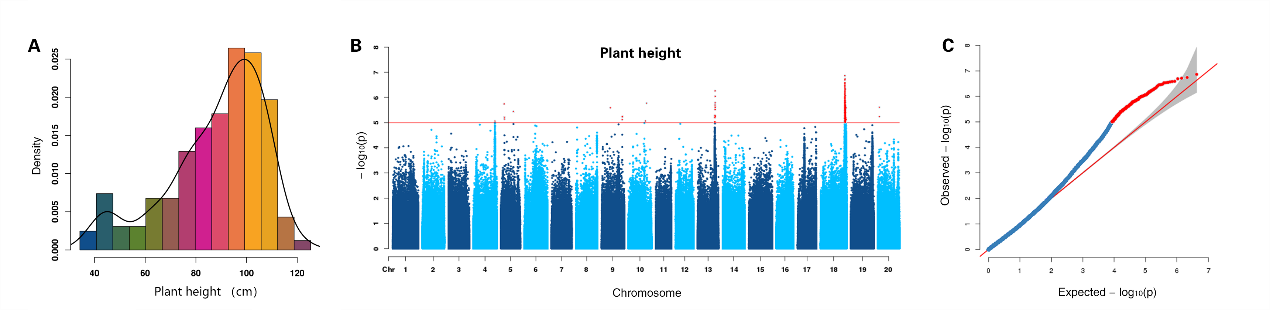

**Figure S14** GWAS of plant height using MLM. **A.** Density distribution of plant height. **B.** Manhattan plots for plant height. Negative log_10_ P-values from a genome-wide scan are plotted against SNP positions of 20 chromosomes. **C.** Quantile-quantile plot for plant height. The horizontal red line indicates the significant threshold (10^-5^). Trait-associated SNPs above the significant threshold are colored in red.

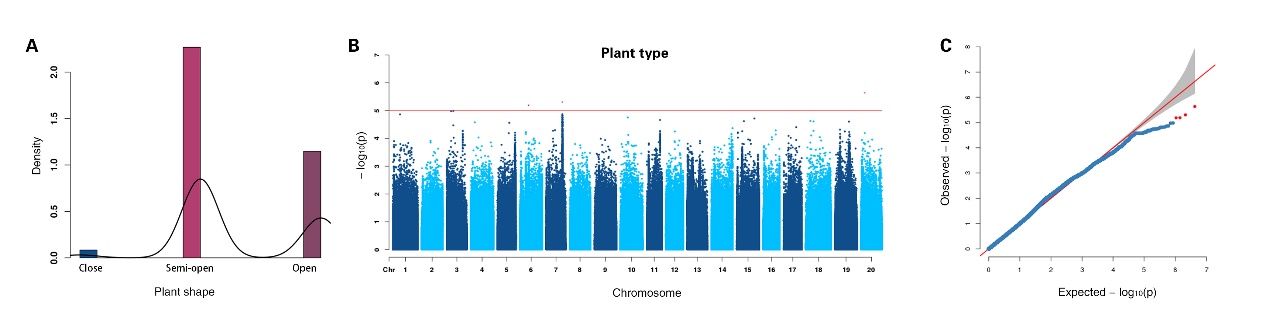

**Figure S15** GWAS of plant type using MLM. **A.** Density distribution of plant type. **B.** Manhattan plots for plant type. Negative log_10_ P-values from a genome-wide scan are plotted against SNP positions of 20 chromosomes. **C.** Quantile-quantile plot for plant type. The horizontal red line indicates the significant threshold (10^-5^). Trait-associated SNPs above the significant threshold are colored in red.

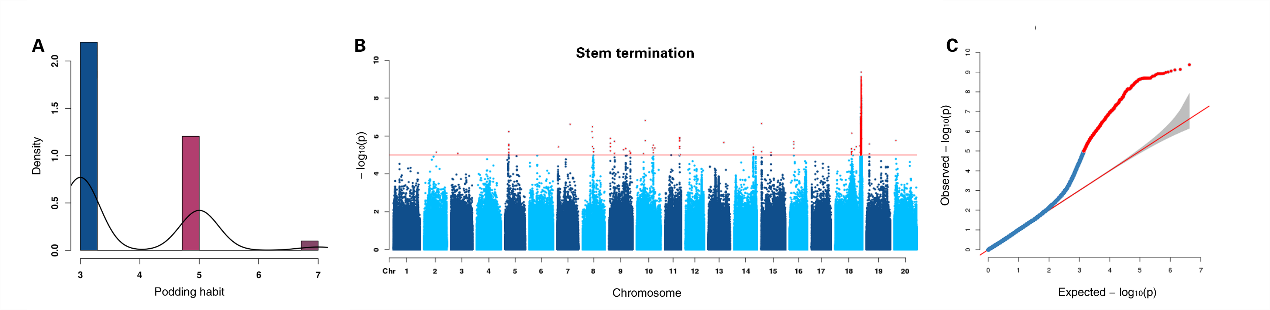

**Figure S16** GWAS of stem termination using MLM. **A.** Density distribution of stem termination. **B.** Manhattan plots for stem termination. Negative log_10_ P-values from a genome-wide scan are plotted against SNP positions of 20 chromosomes. **C.** Quantile-quantile plot for stem termination. The horizontal red line indicates the significant threshold (10^-5^). Trait-associated SNPs above the significant threshold are colored in red.

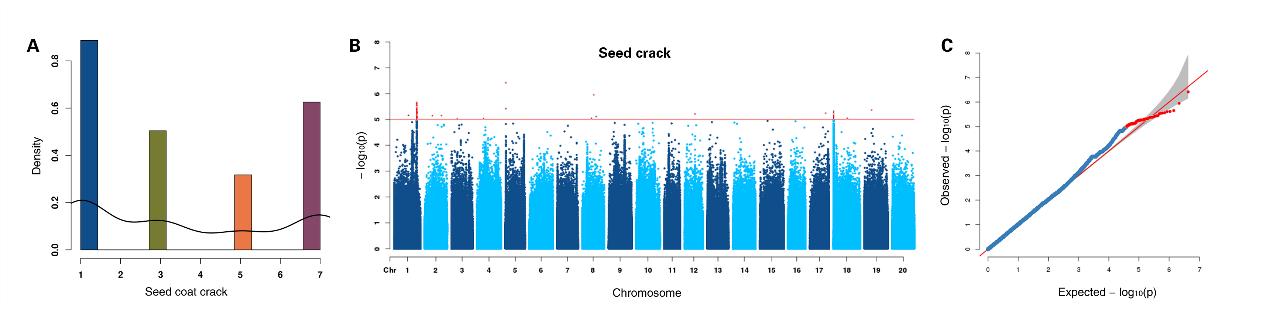

**Figure S17** GWAS of seed crack using MLM. **A.** Density distribution of seed crack. **B.** Manhattan plots for seed crack. Negative log_10_ P-values from a genome-wide scan are plotted against SNP positions of 20 chromosomes. **C.** Quantile-quantile plot for seed crack. The horizontal red line indicates the significant threshold (10^-5^). Trait-associated SNPs above the significant threshold are colored in red.

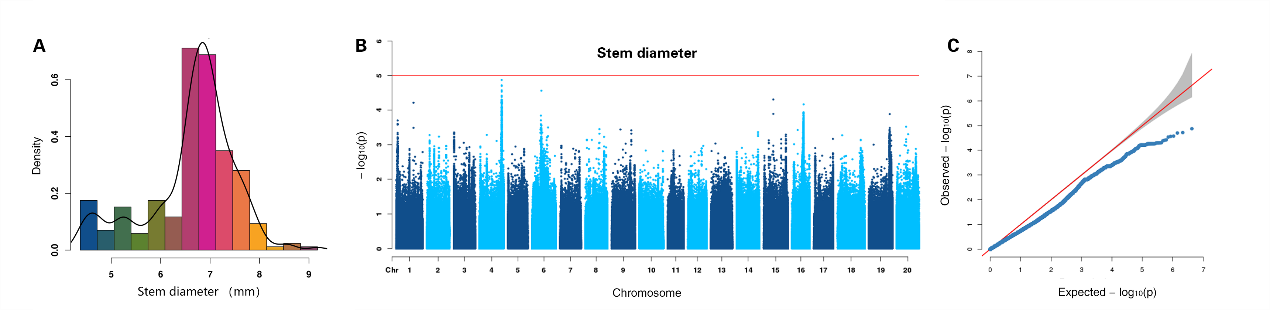

**Figure S18** GWAS of stem diameter using MLM. **A.** Density distribution of stem diameter. **B.** Manhattan plots for stem diameter. Negative log_10_ P-values from a genome-wide scan are plotted against SNP positions of 20 chromosomes. **C.** Quantile-quantile plot for stem diameter. The horizontal red line indicates the significant threshold (10^-5^). Trait-associated SNPs above the significant threshold are colored in red.

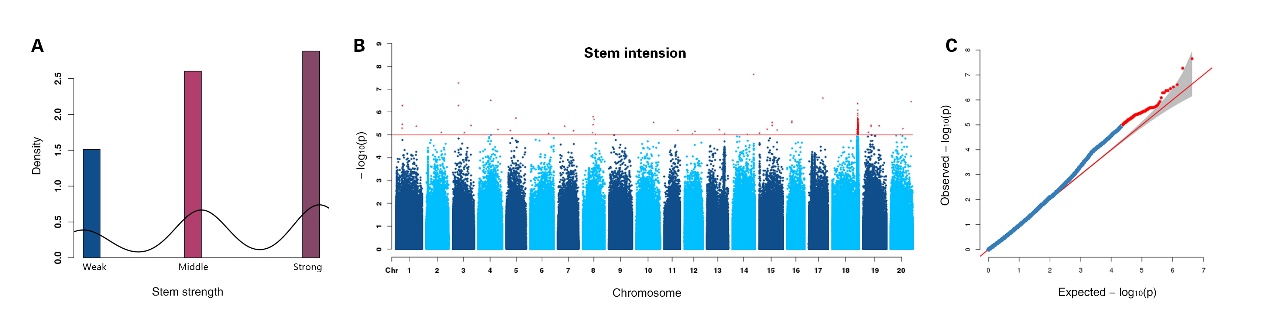

**Figure S19** GWAS of stem intension using MLM. **A.** Density distribution of stem intension. **B.** Manhattan plots for stem intension. Negative log_10_ P-values from a genome-wide scan are plotted against SNP positions of 20 chromosomes. **C.** Quantile-quantile plot for stem intension. The horizontal red line indicates the significant threshold (10^-5^). Trait-associated SNPs above the significant threshold are colored in red.

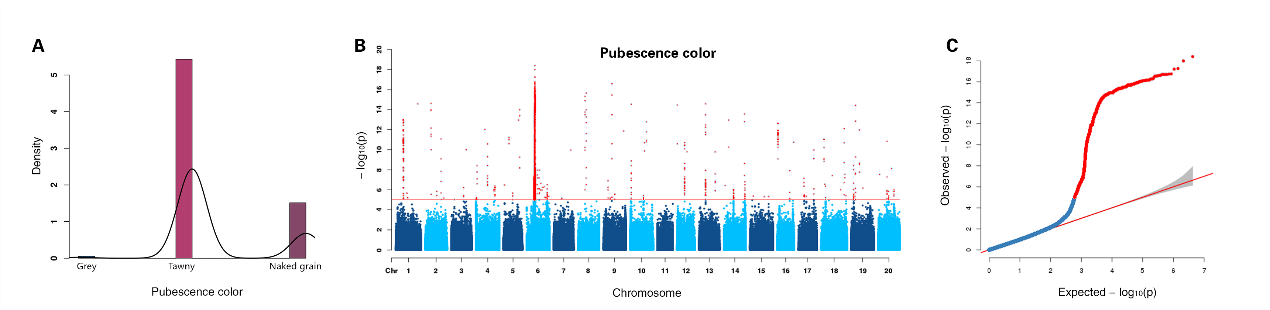

**Figure S20** GWAS of pubescence color using MLM. **A.** Density distribution of pubescence color. **B.** Manhattan plots for pubescence color. Negative log_10_ P-values from a genome-wide scan are plotted against SNP positions of 20 chromosomes. **C.** Quantile-quantile plot for pubescence color. The horizontal red line indicates the significant threshold (10^-5^). Trait-associated SNPs above the significant threshold are colored in red.

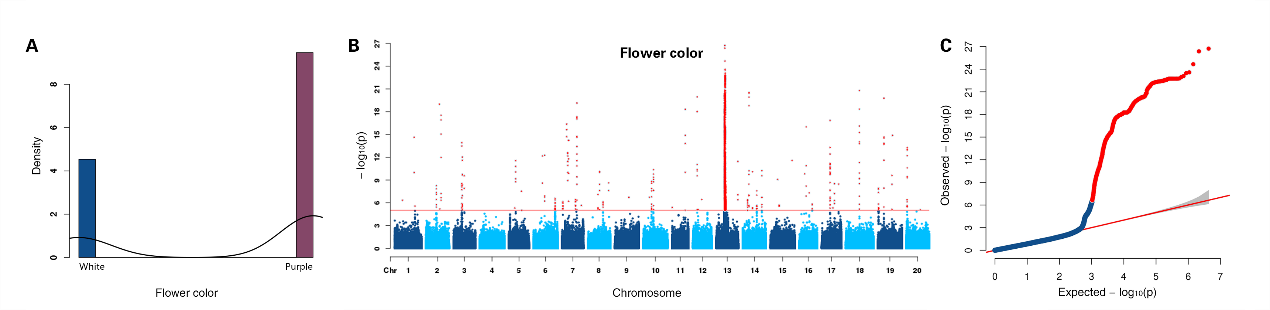

**Figure S21** GWAS of flower color using MLM. **A.** Density distribution of flower color. **B.** Manhattan plots for flower color. Negative log_10_ P-values from a genome-wide scan are plotted against SNP positions of 20 chromosomes. **C.** Quantile-quantile plot for flower color. The horizontal red line indicates the significant threshold (10^-5^). Trait-associated SNPs above the significant threshold are colored in red.

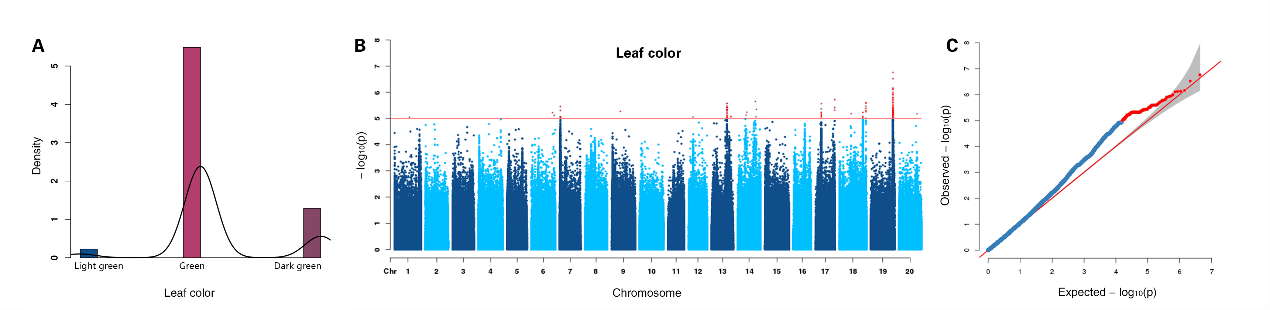

**Figure S22** GWAS of leaf color using MLM. **A.** Density distribution of leaf color. **B.** Manhattan plots for leaf color. Negative log_10_ P-values from a genome-wide scan are plotted against SNP positions of 20 chromosomes. **C.** Quantile-quantile plot for leaf color. The horizontal red line indicates the significant threshold (10^-5^). Trait-associated SNPs above the significant threshold are colored in red.

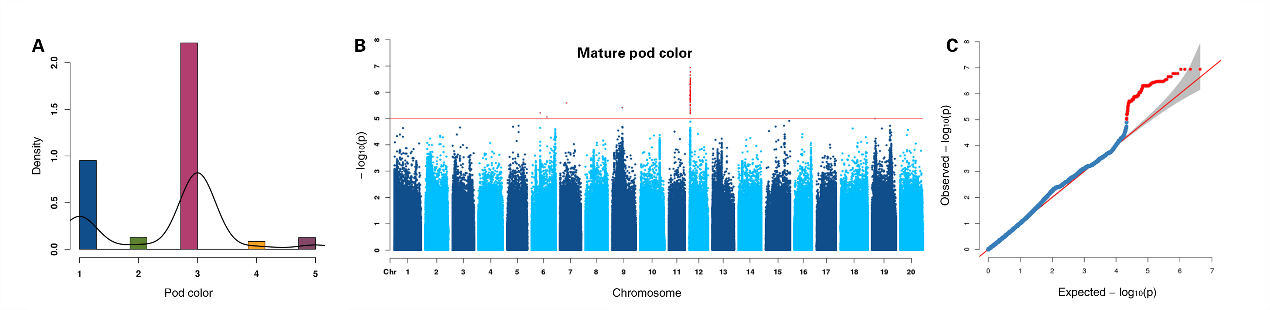

**Figure S23** GWAS of mature pod color using MLM. **A.** Density distribution of mature pod color. **B.** Manhattan plots for mature pod color. Negative log_10_ P-values from a genome-wide scan are plotted against SNP positions of 20 chromosomes. **C.** Quantile-quantile plot for mature pod color. The horizontal red line indicates the significant threshold (10^-5^). Trait-associated SNPs above the significant threshold are colored in red.

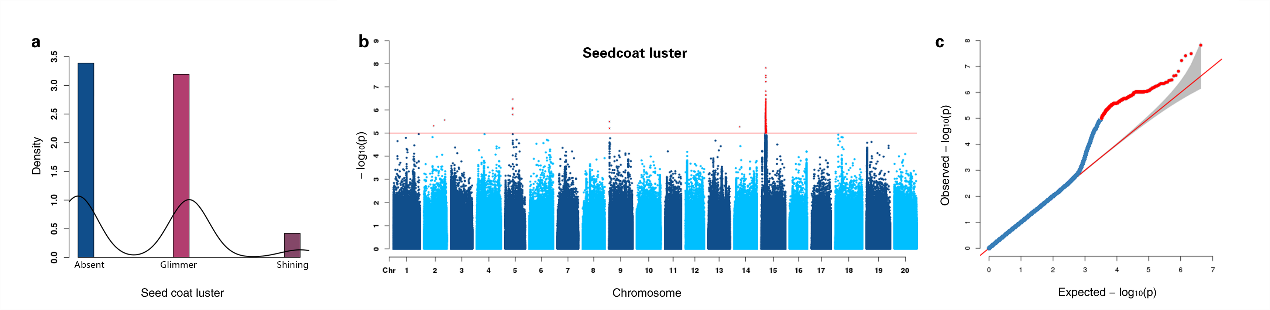

**Figure S24** GWAS of seedcoat luster using MLM. **A.** Density distribution of seedcoat luster. **B.** Manhattan plots for seedcoat luster. Negative log_10_ P-values from a genome-wide scan are plotted against SNP positions of 20 chromosomes. **C.** Quantile-quantile plot for seedcoat luster. The horizontal red line indicates the significant threshold (10^-5^). Trait-associated SNPs above the significant threshold are colored in red.

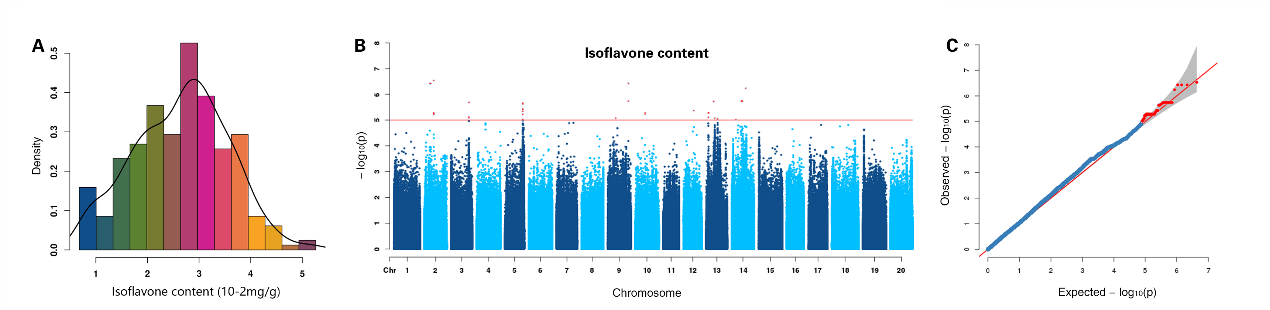

**Figure S25** GWAS of isoflavone content using MLM. **A.** Density distribution of isoflavone content. **B.** Manhattan plots for isoflavone content. Negative log_10_ P-values from a genome-wide scan are plotted against SNP positions of 20 chromosomes. **C.** Quantile-quantile plot for isoflavone content. The horizontal red line indicates the significant threshold (10^-5^). Trait-associated SNPs above the significant threshold are colored in red.

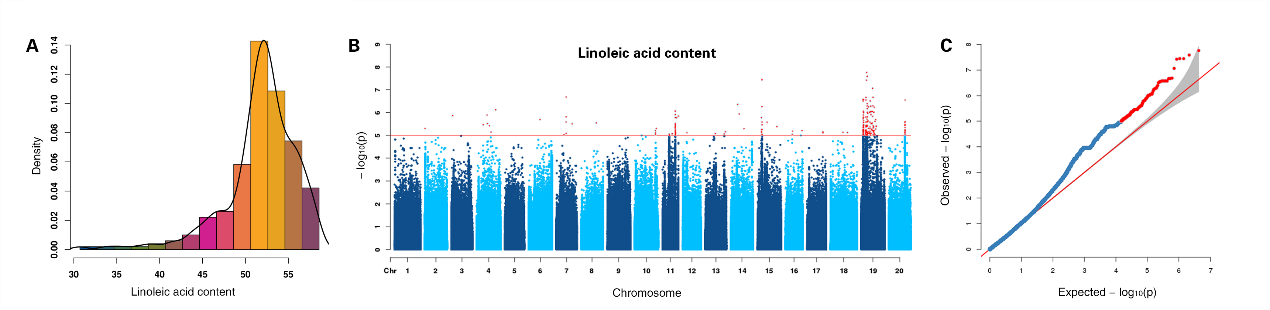

**Figure S26** GWAS of linoleic acid content using MLM. **A.** Density distribution of linoleic acid content. **B.** Manhattan plots for linoleic acid content. Negative log_10_ P-values from a genome-wide scan are plotted against SNP positions of 20 chromosomes. **C.** Quantile-quantile plot for linoleic acid content. The horizontal red line indicates the significant threshold (10^-5^). Trait-associated SNPs above the significant threshold are colored in red.

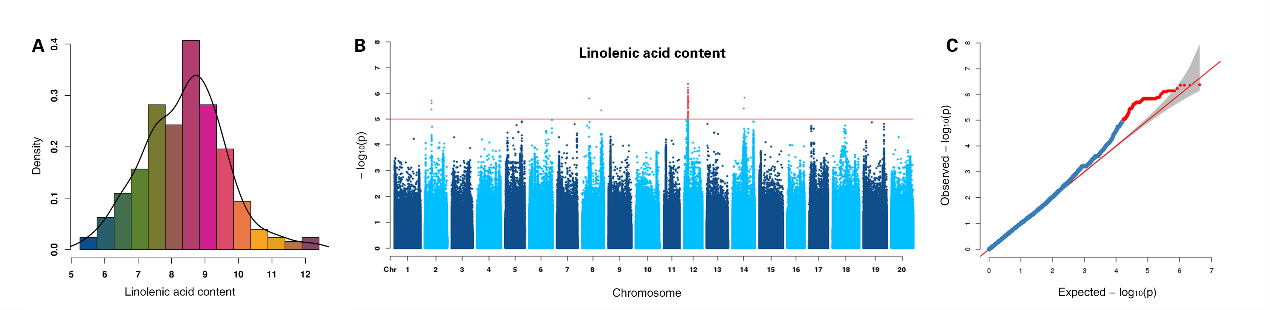

**Figure S27** GWAS of linolenic acid content using MLM. **A.** Density distribution of linolenic acid content. **B.** Manhattan plots for linolenic acid content. Negative log_10_ P-values from a genome-wide scan are plotted against SNP positions of 20 chromosomes. **C.** Quantile-quantile plot for linolenic acid content. The horizontal red line indicates the significant threshold (10^-5^). Trait-associated SNPs above the significant threshold are colored in red.

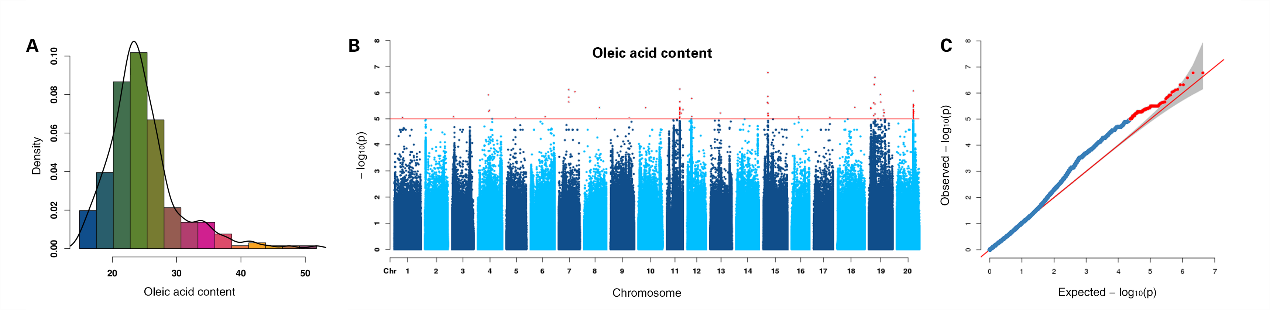

**Figure S28** GWAS of oleic acid content using MLM. **A.** Density distribution of oleic acid content. **B.** Manhattan plots for oleic acid content. Negative log_10_ P-values from a genome-wide scan are plotted against SNP positions of 20 chromosomes. **C.** Quantile-quantile plot for oleic acid content. The horizontal red line indicates the significant threshold (10^-5^). Trait-associated SNPs above the significant threshold are colored in red.

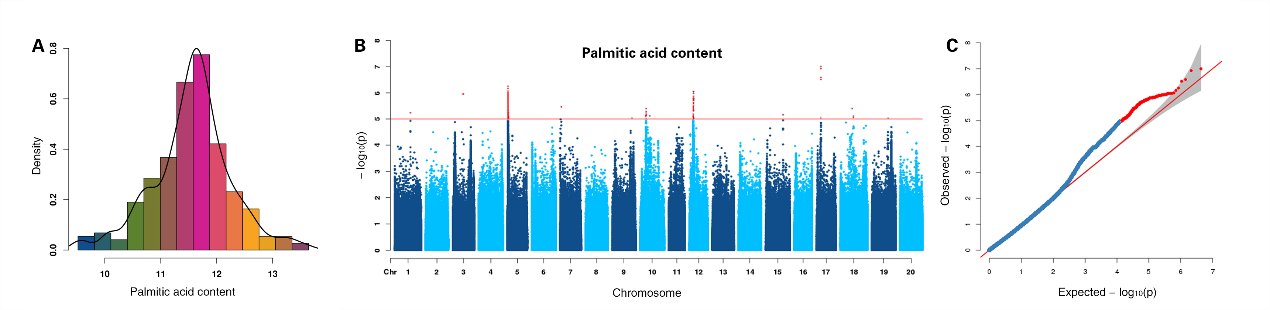

**Figure S29** GWAS of palmitic acid content using MLM. **A.** Density distribution of palmitic acid content. **B.** Manhattan plots for palmitic acid content. Negative log_10_ P-values from a genome-wide scan are plotted against SNP positions of 20 chromosomes. **C.** Quantile-quantile plot for palmitic acid content. The horizontal red line indicates the significant threshold (10^-5^). Trait-associated SNPs above the significant threshold are colored in red.

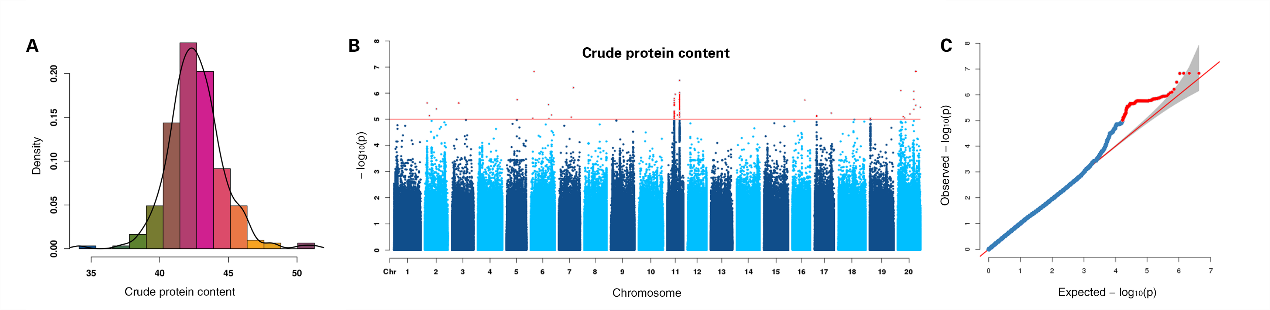

**Figure S30** GWAS of crude protein content using MLM. **A.** Density distribution of crude protein content. **B.** Manhattan plots for crude protein content. Negative log_10_ P-values from a genome-wide scan are plotted against SNP positions of 20 chromosomes. **C.** Quantile-quantile plot for crude protein content. The horizontal red line indicates the significant threshold (10^-5^). Trait-associated SNPs above the significant threshold are colored in red.

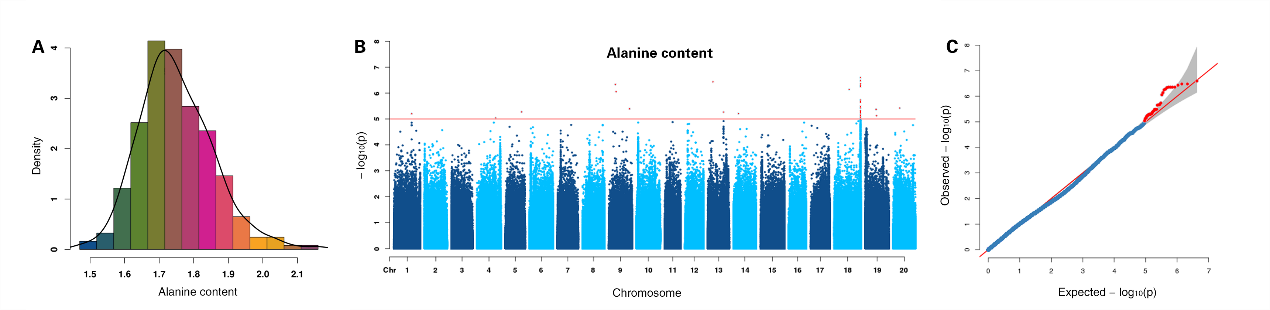

**Figure S31** GWAS of alanine content using MLM. **A.** Density distribution of alanine content. **B.** Manhattan plots for alanine content. Negative log_10_ P-values from a genome-wide scan are plotted against SNP positions of 20 chromosomes. **C.** Quantile-quantile plot for alanine content. The horizontal red line indicates the significant threshold (10^-5^). Trait-associated SNPs above the significant threshold are colored in red.

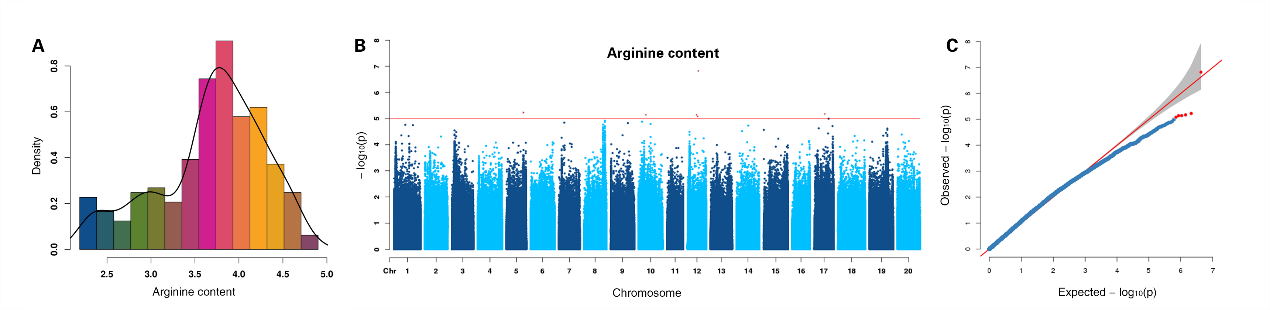

**Figure S32** GWAS of arginine content using MLM. **A.** Density distribution of arginine content. **B.** Manhattan plots for arginine content. Negative log_10_ P-values from a genome-wide scan are plotted against SNP positions of 20 chromosomes. **C.** Quantile-quantile plot for arginine content. The horizontal red line indicates the significant threshold (10^-5^). Trait-associated SNPs above the significant threshold are colored in red.

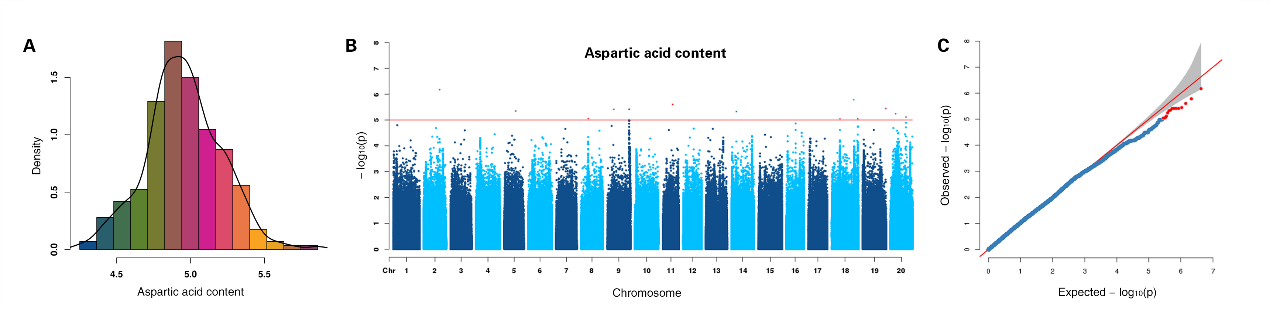

**Figure S33** GWAS of aspartic acid content using MLM. **A.** Density distribution of aspartic acid content. **B.** Manhattan plots for aspartic acid content. Negative log_10_ P-values from a genome-wide scan are plotted against SNP positions of 20 chromosomes. **C.** Quantile-quantile plot for aspartic acid content. The horizontal red line indicates the significant threshold (10^-5^). Trait-associated SNPs above the significant threshold are colored in red.

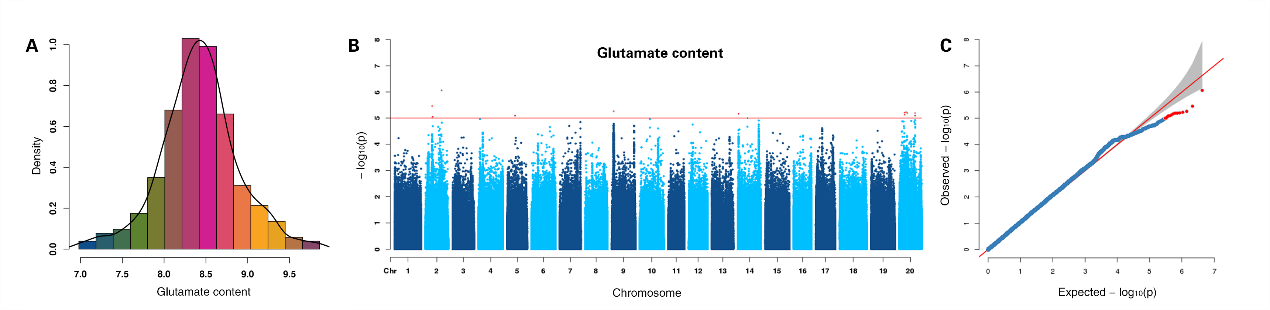

**Figure S34** GWAS of glutamate content using MLM. **A.** Density distribution of glutamate content. **B.** Manhattan plots for glutamate content. Negative log_10_ P-values from a genome-wide scan are plotted against SNP positions of 20 chromosomes. **C.** Quantile-quantile plot for glutamate content. The horizontal red line indicates the significant threshold (10^-5^). Trait-associated SNPs above the significant threshold are colored in red.

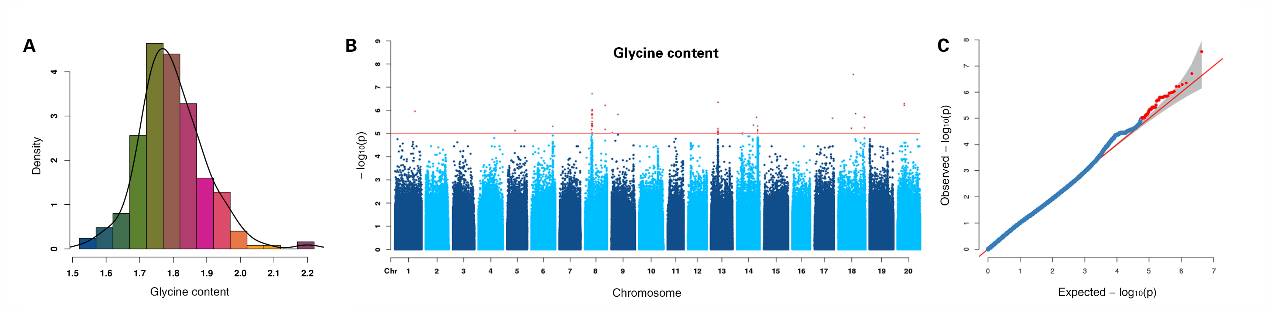

**Figure S35** GWAS of glycine content using MLM. **A.** Density distribution of glycine content. **B.** Manhattan plots for glycine content. Negative log_10_ P-values from a genome-wide scan are plotted against SNP positions of 20 chromosomes. **C.** Quantile-quantile plot for glycine content. The horizontal red line indicates the significant threshold (10^-5^). Trait-associated SNPs above the significant threshold are colored in red.

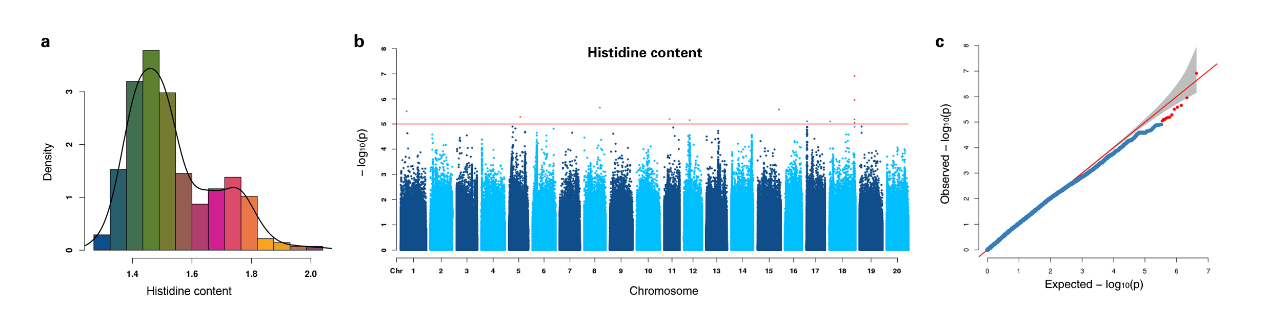

**Figure S36** GWAS of histidine content using MLM. **A.** Density distribution of histidine content. **B.** Manhattan plots for histidine content. Negative log_10_ P-values from a genome-wide scan are plotted against SNP positions of 20 chromosomes. **C.** Quantile-quantile plot for histidine content. The horizontal red line indicates the significant threshold (10^-5^). Trait-associated SNPs above the significant threshold are colored in red.

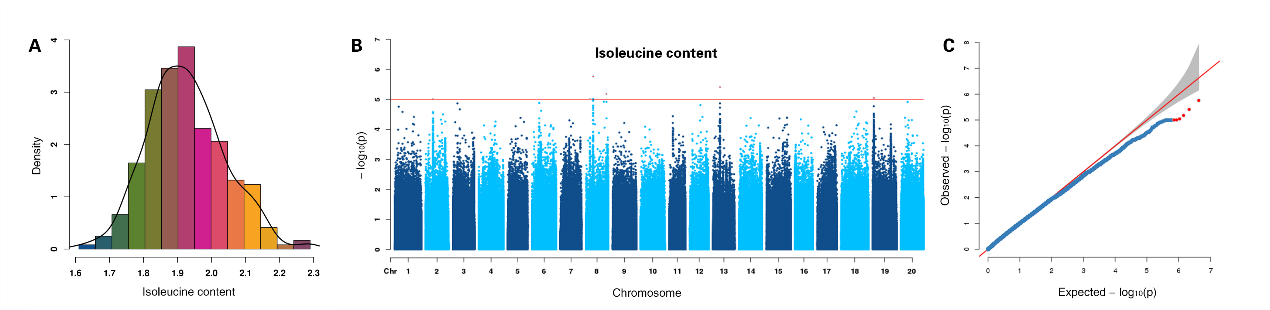

**Figure S37** GWAS of isoleucine content using MLM. **A.** Density distribution of isoleucine content. **B.** Manhattan plots for isoleucine content. Negative log_10_ P-values from a genome-wide scan are plotted against SNP positions of 20 chromosomes. **C.** Quantile-quantile plot for isoleucine content. The horizontal red line indicates the significant threshold (10^-5^). Trait-associated SNPs above the significant threshold are colored in red.

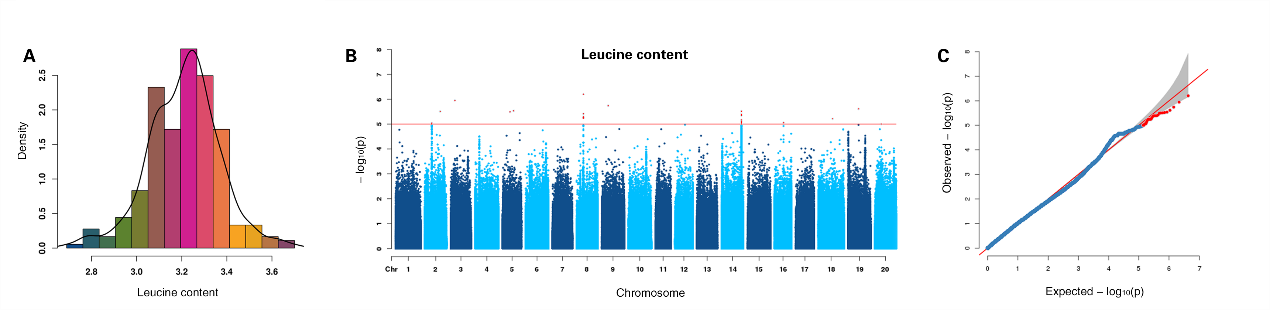

**Figure S38** GWAS of leucine content using MLM. **A.** Density distribution of leucine content. **B.** Manhattan plots for leucine content. Negative log_10_ P-values from a genome-wide scan are plotted against SNP positions of 20 chromosomes. **C.** Quantile-quantile plot for leucine content. The horizontal red line indicates the significant threshold (10^-5^). Trait-associated SNPs above the significant threshold are colored in red.

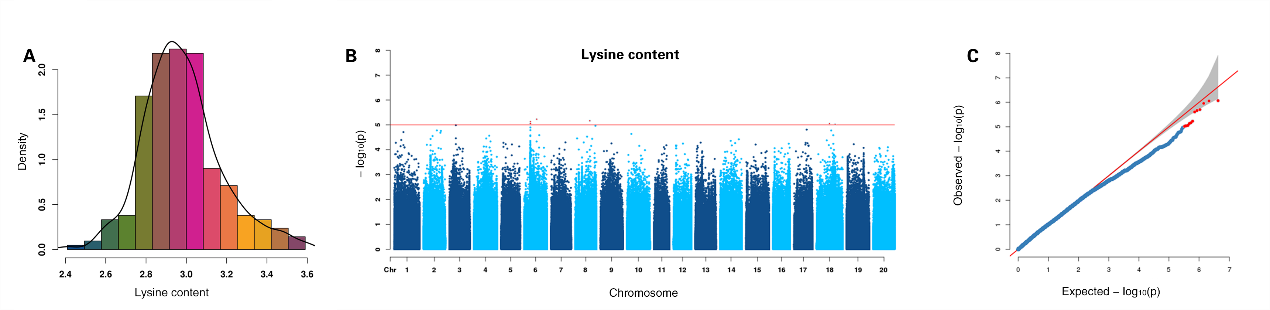

**Figure S39** GWAS of lysine content using MLM. **A.** Density distribution of lysine content. **B.** Manhattan plots for lysine content. Negative log_10_ P-values from a genome-wide scan are plotted against SNP positions of 20 chromosomes. **C.** Quantile-quantile plot for lysine content. The horizontal red line indicates the significant threshold (10^-5^). Trait-associated SNPs above the significant threshold are colored in red.

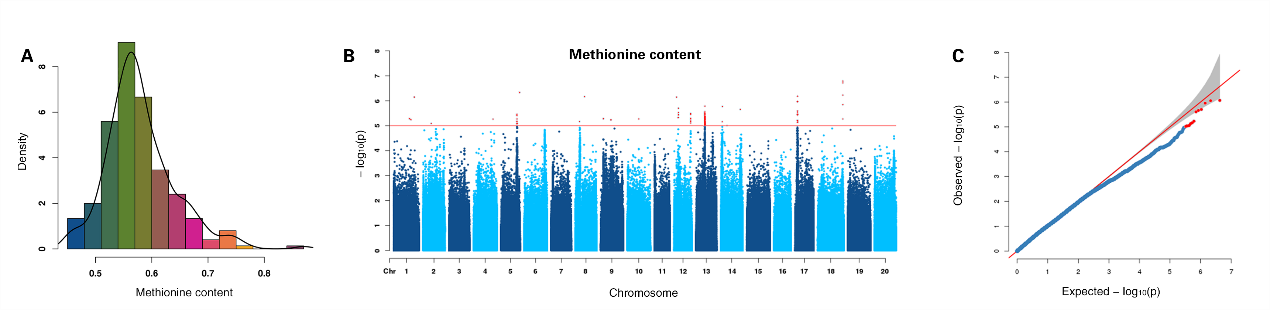

**Figure S40** GWAS of methionine content using MLM. **A.** Density distribution of methionine content. **B.** Manhattan plots for methionine content. Negative log_10_ P-values from a genome-wide scan are plotted against SNP positions of 20 chromosomes. **C.** Quantile-quantile plot for methionine content. The horizontal red line indicates the significant threshold (10^-5^). Trait-associated SNPs above the significant threshold are colored in red.

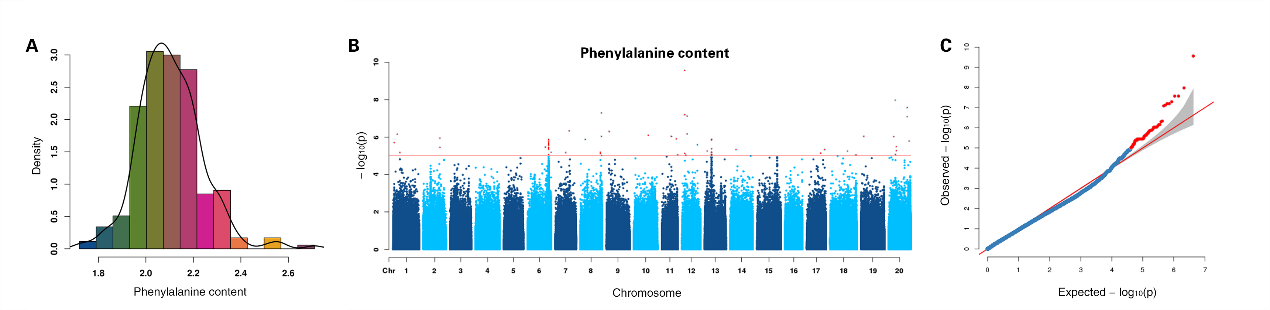

**Figure S41** GWAS of phenylalanine content using MLM. **A.** Density distribution of phenylalanine content. **B.** Manhattan plots for phenylalanine content. Negative log_10_ P-values from a genome-wide scan are plotted against SNP positions of 20 chromosomes. **C.** Quantile-quantile plot for phenylalanine content. The horizontal red line indicates the significant threshold (10^-5^). Trait-associated SNPs above the significant threshold are colored in red.

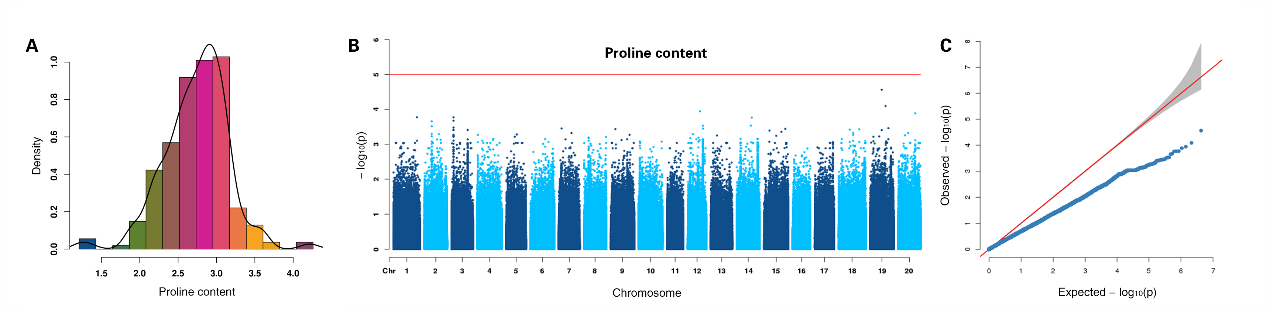

**Figure S42** GWAS of proline content using MLM. **A.** Density distribution of proline content. **B.** Manhattan plots for proline content. Negative log_10_ P-values from a genome-wide scan are plotted against SNP positions of 20 chromosomes. **C.** Quantile-quantile plot for proline content. The horizontal red line indicates the significant threshold (10^-5^). Trait-associated SNPs above the significant threshold are colored in red.

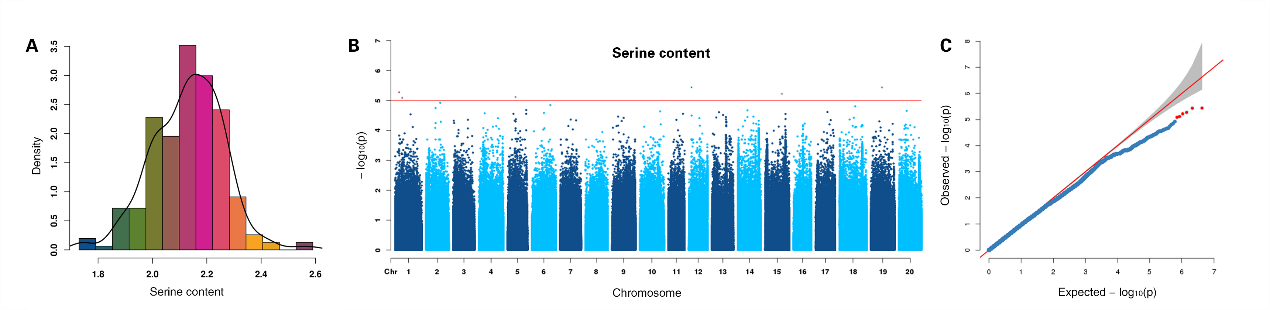

**Figure S43** GWAS of serine content using MLM. **A.** Density distribution of serine content. **B.** Manhattan plots for serine content. Negative log_10_ P-values from a genome-wide scan are plotted against SNP positions of 20 chromosomes. **C.** Quantile-quantile plot for serine content. The horizontal red line indicates the significant threshold (10^-5^). Trait-associated SNPs above the significant threshold are colored in red.

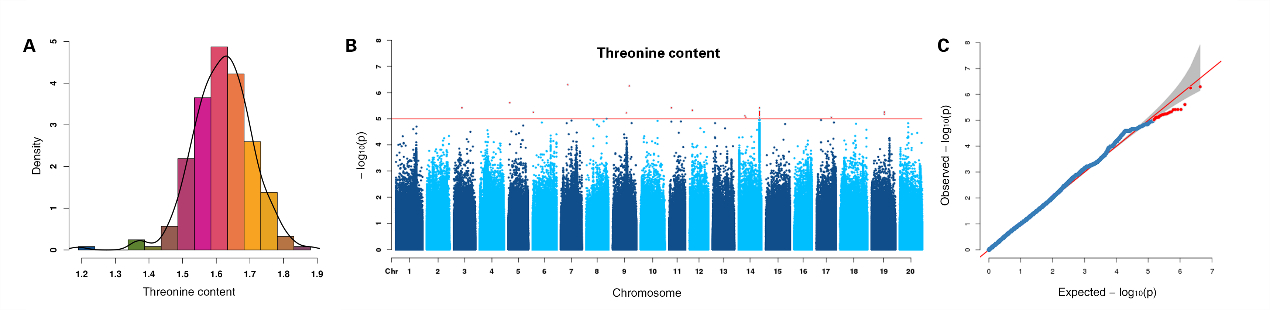

**Figure S44** GWAS of threonine content using MLM. **A.** Density distribution of threonine content. **B.** Manhattan plots for threonine content. Negative log_10_ P-values from a genome-wide scan are plotted against SNP positions of 20 chromosomes. **C.** Quantile-quantile plot for threonine content. The horizontal red line indicates the significant threshold (10^-5^). Trait-associated SNPs above the significant threshold are colored in red.

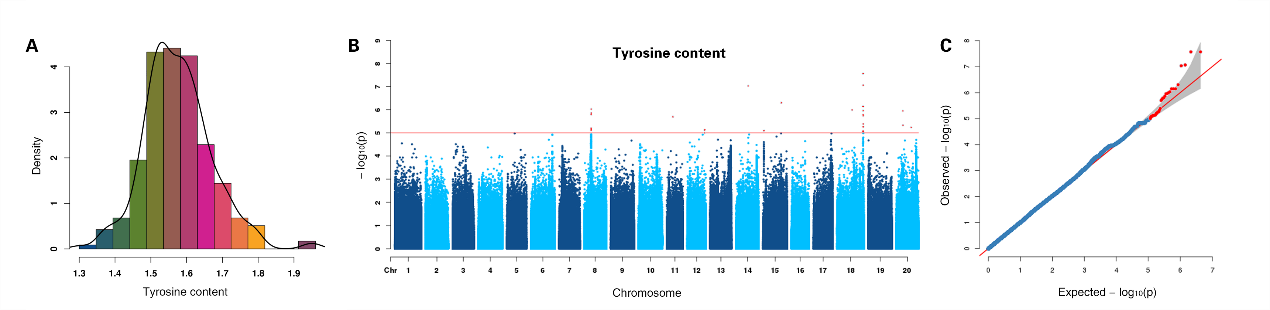

**Figure S45** GWAS of tyrosine content using MLM. **A.** Density distribution of tyrosine content. **B.** Manhattan plots for tyrosine content. Negative log_10_ P-values from a genome-wide scan are plotted against SNP positions of 20 chromosomes. **C.** Quantile-quantile plot for tyrosine content. The horizontal red line indicates the significant threshold (10^-5^). Trait-associated SNPs above the significant threshold are colored in red.

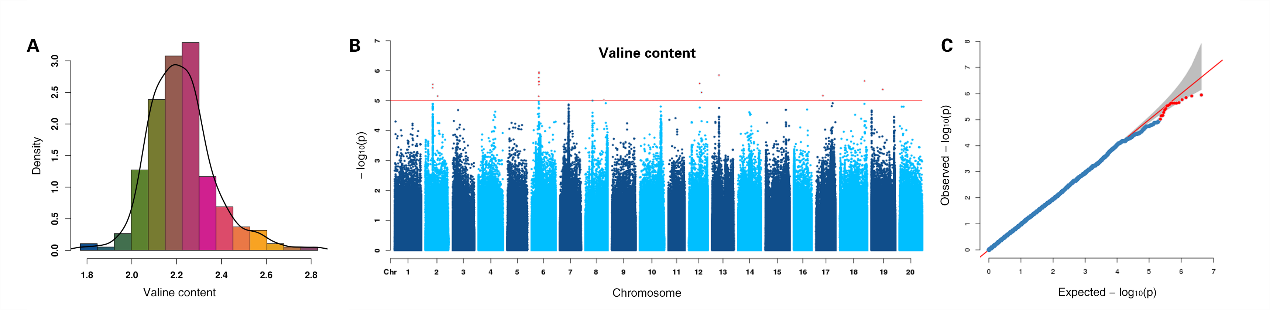

**Figure S46** GWAS of valine content using MLM. **A.** Density distribution of valine content. **B.** Manhattan plots for valine content. Negative log_10_ P-values from a genome-wide scan are plotted against SNP positions of 20 chromosomes. **C.** Quantile-quantile plot for valine content. The horizontal red line indicates the significant threshold (10^-5^). Trait-associated SNPs above the significant threshold are colored in red.

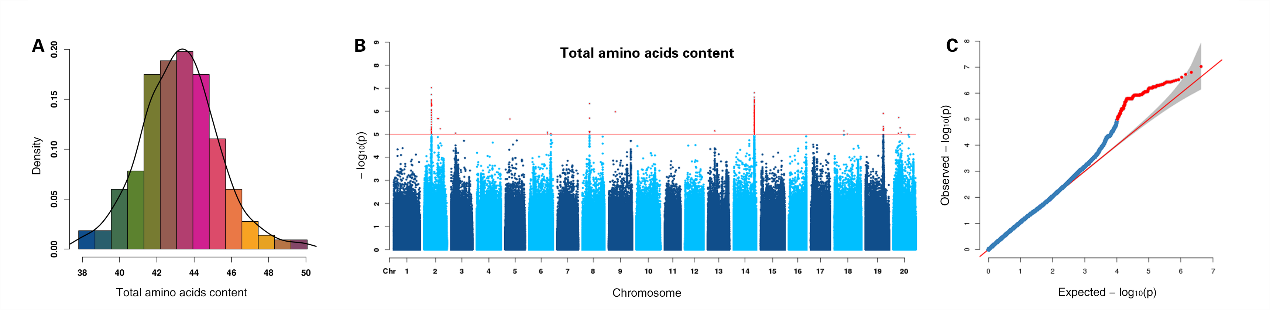

**Figure S47** GWAS of total amino acids content using MLM. **A.** Density distribution of total amino acids content. **B.** Manhattan plots for total amino acids content. Negative log_10_ P-values from a genome-wide scan are plotted against SNP positions of 20 chromosomes. **C.** Quantile-quantile plot for total amino acids content. The horizontal red line indicates the significant threshold (10^-5^). Trait-associated SNPs above the significant threshold are colored in red.

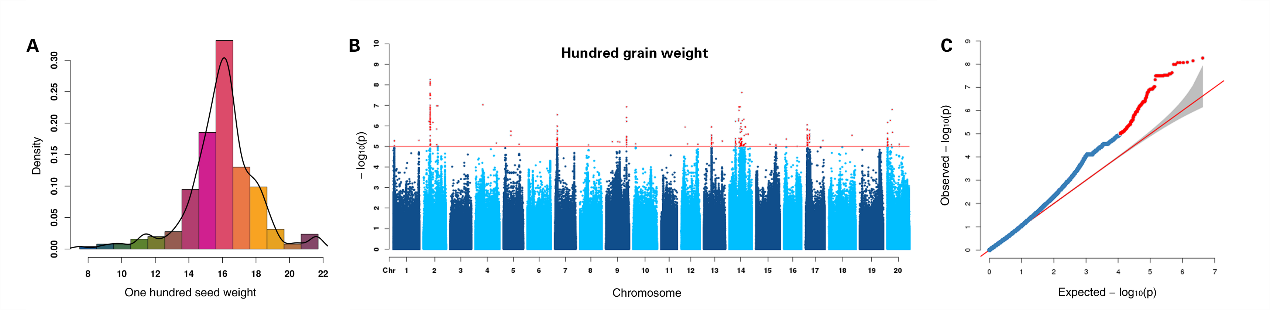

**Figure S48** GWAS of hundred grain weight using MLM. **A.** Density distribution of hundred grain weight. **B.** Manhattan plots for hundred grain weight. Negative log_10_ P-values from a genome-wide scan are plotted against SNP positions of 20 chromosomes. **C.** Quantile-quantile plot for hundred grain weight. The horizontal red line indicates the significant threshold (10^-5^). Trait-associated SNPs above the significant threshold are colored in red.

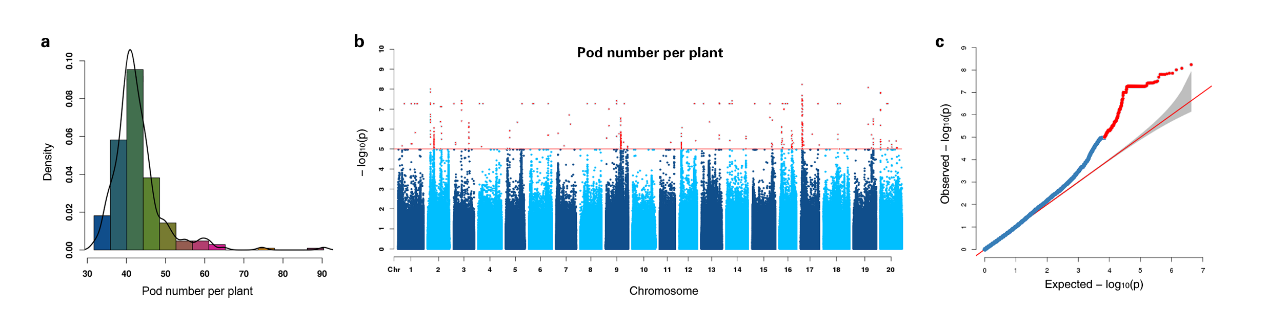

**Figure S49** GWAS of pod number per plant using MLM. **A.** Density distribution of pod number per plant. **B.** Manhattan plots for pod number per plant. Negative log_10_ P-values from a genome-wide scan are plotted against SNP positions of 20 chromosomes. **C.** Quantile-quantile plot for pod number per plant. The horizontal red line indicates the significant threshold (10^-5^). Trait-associated SNPs above the significant threshold are colored in red.

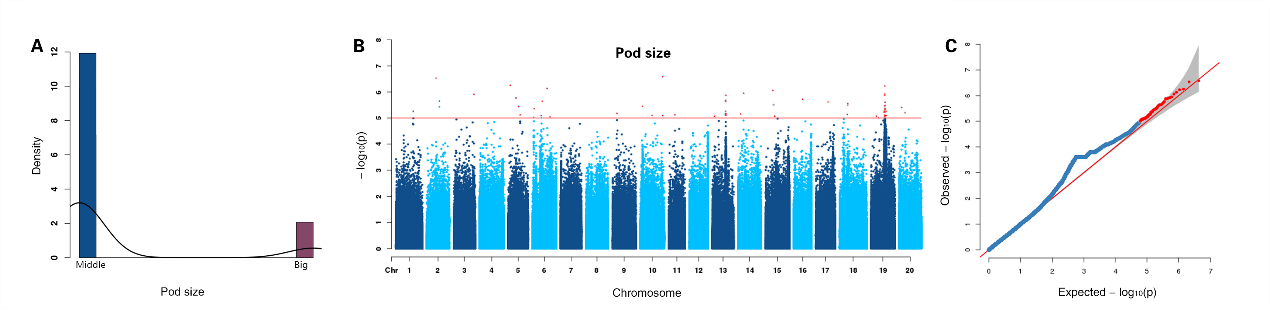

**Figure S50** GWAS of pod size using MLM. **A.** Density distribution of pod size. **B.** Manhattan plots for pod size. Negative log_10_ P-values from a genome-wide scan are plotted against SNP positions of 20 chromosomes. **C.** Quantile-quantile plot for pod size. The horizontal red line indicates the significant threshold (10^-5^). Trait-associated SNPs above the significant threshold are colored in red.

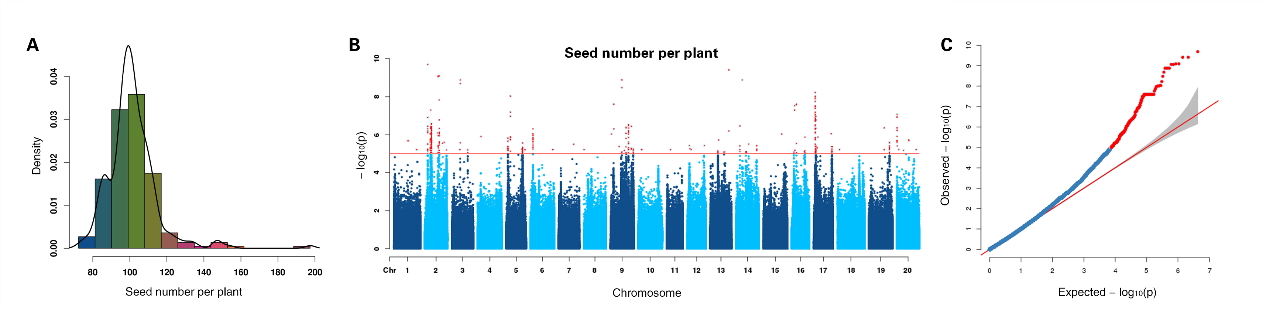

**Figure S51** GWAS of seed number per plant using MLM. **A.** Density distribution of seed number per plant. **B.** Manhattan plots for seed number per plant. Negative log_10_ P-values from a genome-wide scan are plotted against SNP positions of 20 chromosomes. **C.** Quantile-quantile plot for seed number per plant. The horizontal red line indicates the significant threshold (10^-5^). Trait-associated SNPs above the significant threshold are colored in red.

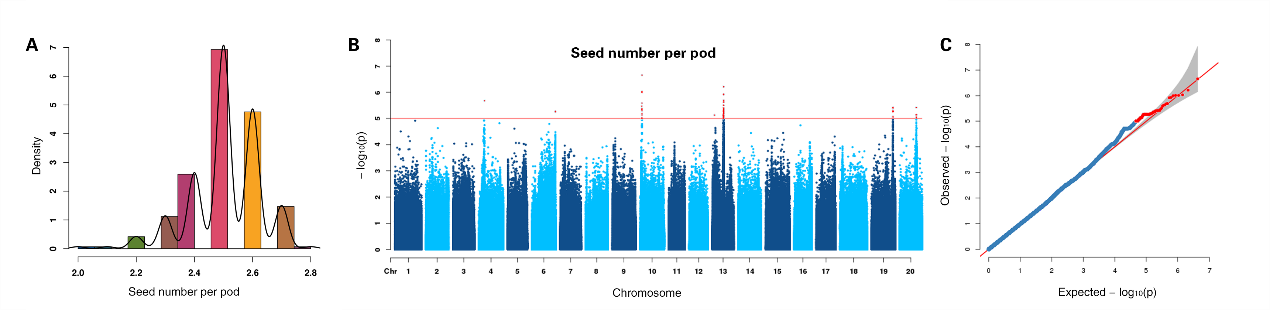

**Figure S52** GWAS of seed number per pod using MLM. **A.** Density distribution of seed number per pod. **B.** Manhattan plots for seed number per pod. Negative log_10_ P-values from a genome-wide scan are plotted against SNP positions of 20 chromosomes. **C.** Quantile-quantile plot for seed number per pod. The horizontal red line indicates the significant threshold (10^-5^). Trait-associated SNPs above the significant threshold are colored in red.

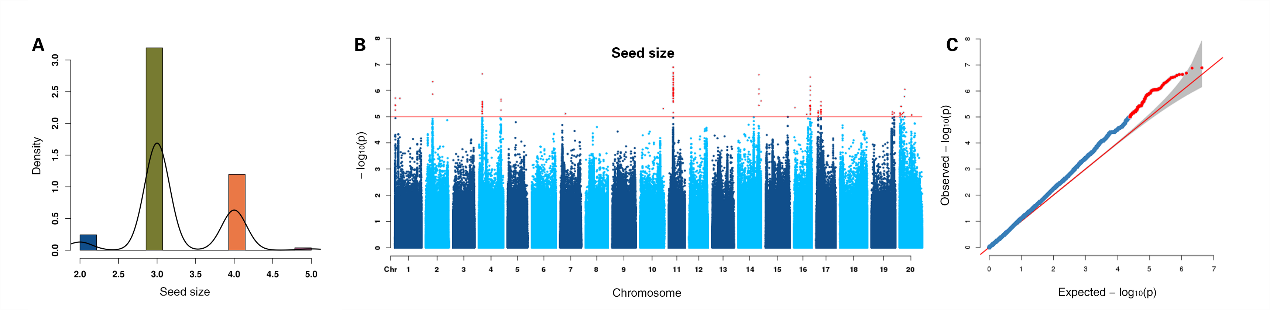

**Figure S53** GWAS of seed size using MLM. **A.** Density distribution of seed size. **B.** Manhattan plots for seed size. Negative log_10_ P-values from a genome-wide scan are plotted against SNP positions of 20 chromosomes. **C.** Quantile-quantile plot for seed size. The horizontal red line indicates the significant threshold (10^-5^). Trait-associated SNPs above the significant threshold are colored in red.

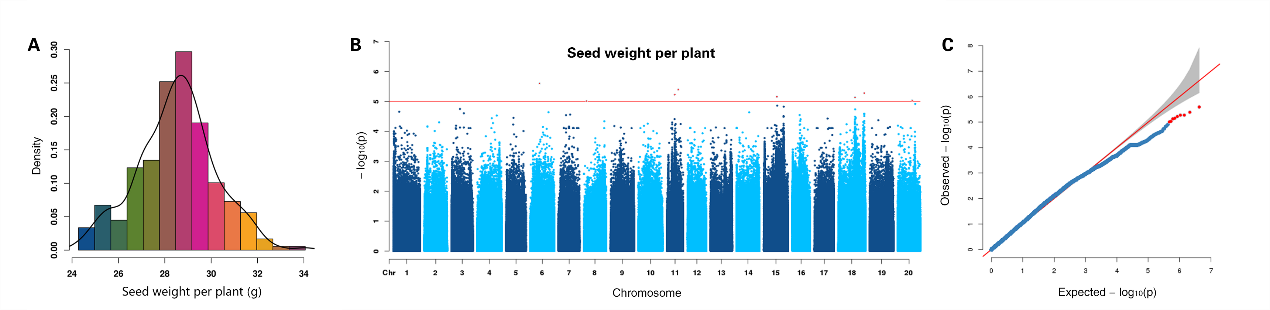

**Figure S54** GWAS of seed weight per plant using MLM. **A.** Density distribution of seed weight per plant. **B.** Manhattan plots for seed weight per plant. Negative log_10_ P-values from a genome-wide scan are plotted against SNP positions of 20 chromosomes. **C.** Quantile-quantile plot for seed weight per plant. The horizontal red line indicates the significant threshold (10^-5^). Trait-associated SNPs above the significant threshold are colored in red.

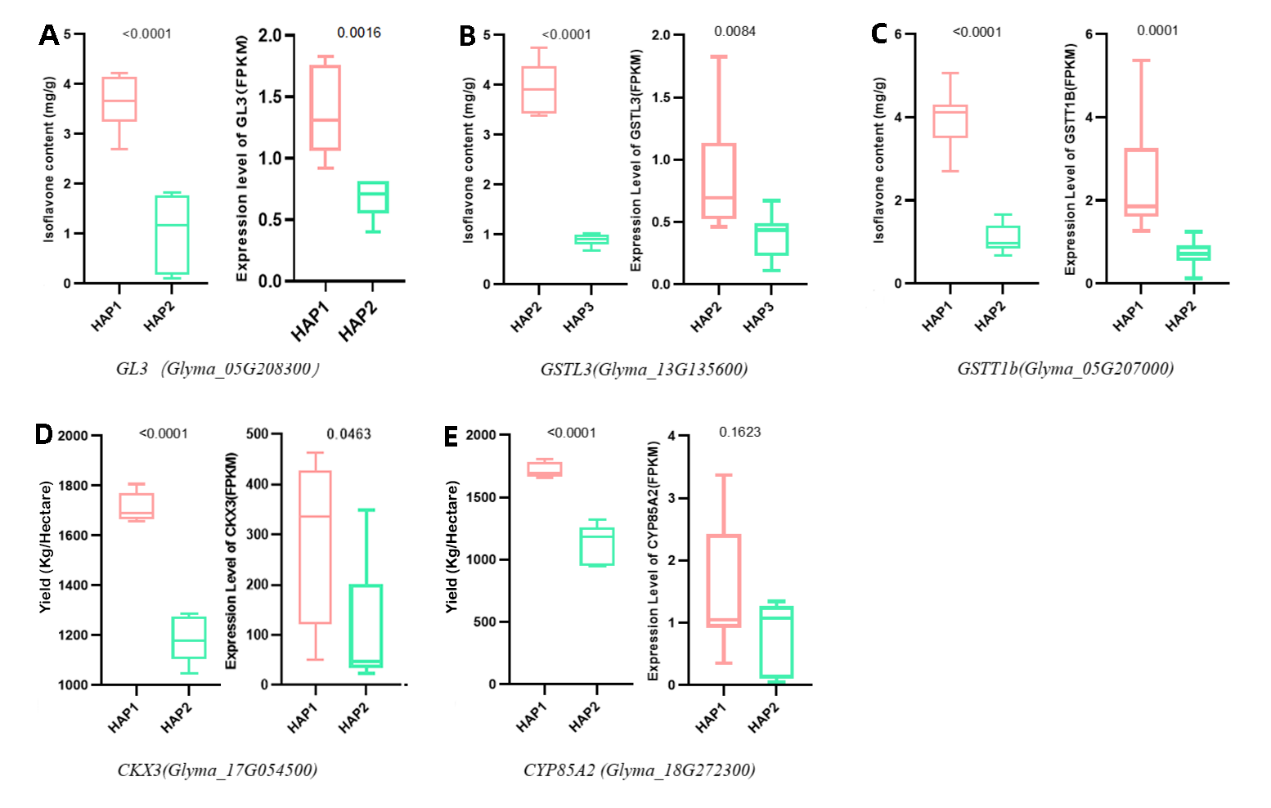

**Figure S55** Gene expression validation of different haplotypes/phenotypes for five candidate genes. **A.** Phenotype distribution (left) and expression level (right) of *GL3* for different haplotypes. **B.** Phenotype distribution (left) and expression level (right) of *GSTL3* for different haplotypes. **C.** Phenotype distribution (left) and expression level (right) of *GSTT1b* for different haplotypes. **D.** Phenotype distribution (left) and expression level (right) of *CKX3* for different haplotypes. **E.** Phenotype distribution (left) and expression level (right) of *CYP85A2* for different haplotypes.
